# Impact of *Bmal1* rescue and time-restricted feeding on liver and muscle proteomes during the active phase in mice

**DOI:** 10.1101/2023.06.12.544652

**Authors:** Jacob G. Smith, Jeffrey Molendijk, Ronnie Blazev, Wan Hsi Chen, Qing Zhang, Christopher Litwin, Carolina M. Greco, Paolo Sassone-Corsi, Pura Muñoz-Cánoves, Benjamin L. Parker, Kevin B. Koronowski

## Abstract

**Objective:** Molecular clocks and daily feeding cycles support metabolism in peripheral tissues. Although the roles of local clocks and feeding is well defined at the transcriptional level, their impact on governing protein abundances in peripheral tissues is unclear. Here, we determine the relative contributions of the local molecular clock and daily feeding cycles on liver and muscle proteomes during feeding.

**Methods:** LC-MS/MS was performed on liver and skeletal muscle harvested four hours into the dark phase from wild-type (WT), *Bmal1* knockout (KO), and liver- and muscle-*Bmal1*-rescued (LMRE) mice housed under 12-h light/12-h dark cycles with either *ad libitum* feeding or nighttime-restricted feeding. Additional molecular and metabolic analyses were performed on liver and cultured hepatocytes.

**Results:** Feeding-fasting cycles had only minimal effects on liver and none on muscle. In contrast, *Bmal1* KO altered the abundance of 674 proteins in liver, and 80 in muscle. Rescue of liver and muscle *Bmal1* restored 50% of proteins in liver and 25% in muscle. These included proteins involved in carbohydrate metabolism in muscle and in fatty acid oxidation in liver. For liver, proteins involved in *de novo* lipogenesis were largely dependent on *Bmal1* function in other tissues (i.e., the wider clock system). Proteins regulated by BMAL1 were enriched for secreted proteins; we determined that the maintenance of FGF1 abundance requires liver BMAL1, and that autocrine signaling through FGF1 is necessary and sufficient for mitochondrial respiration in hepatocytes.

**Conclusions:** BMAL1 in liver and muscle is a more potent regulator of dark phase proteomes than daily feeding cycles, highlighting the need to assess protein levels in addition to mRNA when investigating clock mechanisms. The proteome is more extensively regulated by BMAL1 in liver than in muscle, and numerous metabolic pathways in peripheral tissues are reliant on the function of the clock system as a whole.

## 1. INTRODUCTION

The mammalian circadian clock system drives 24-hour rhythms of physiology, coordinating metabolic processes within and between tissues. Disruption of the clock system via genetic or environmental means induces metabolic dysregulation and increases risk for metabolic disease [1–3]. The central clock in the suprachiasmatic nucleus regulates systems-level coordination through behavioral cycles, including locomotion and feeding-fasting behavior, which cooperate with humoral and neuronal mechanisms to signal to peripheral tissue clocks. Daily transcriptional programs that support metabolic function are dependent on this coordination, yet how local and central clock signals interact at the proteomic level is unclear.

Two main components of circadian control in peripheral tissues have been described. The local autonomous clock controls a subset of transcriptional rhythms in each tissue, with metabolic function in each tissue similarly restricted [4–6], and the centrally-driven feeding-fasting rhythm engages systemic hormonal signals (e.g., insulin release from the pancreas [5; 7; 8]) and regulates transcription through nutrient responsive signaling pathways [5; 9]. Modulation of feeding-fasting cycles through time-restricted feeding has numerous beneficial effects on metabolism, such as improved insulin sensitivity and reduction of adiposity. Mouse studies show that time-restricted feeding has both clock -dependent and -independent effects [10; 11]. Indeed, our recent work has shown that local clocks and feeding cycles interact to bolster rhythms in liver and muscle, and that peripheral clock function is shaped by interactions with other peripheral clocks [12]. For example, the glucose intolerance of *Bmal1*-null mice was most improved when both liver and muscle clocks were rescued and engaged with feeding cycles. Thus, various levels of clock control and their interactions are important for metabolic homeostasis.

Most studies, including our own, have focused on transcription using mRNA analyses. However, there are many regulatory steps between transcriptional output and protein products, and posttranscriptional mechanisms of clock control are evident [13–15]. Few studies using clock mutant mice, especially *Bmal1*-null mice, have interrogated clock mechanisms at the protein level [16–18], such that much less is known about how the clock system controls the proteomic landscape in each tissue. Here, we address how local clocks and feeding cycles impact metabolic pathways at the protein level in liver and muscle. By considering this critical juncture, we move closer to understanding how time-restricted feeding and modulation of clock function may be targeted for therapeutic gain.

## 2. MATERIAL AND METHODS

### 2.1. Animal experiments

Mice were bred and housed in a vivarium at the University of California Irvine. All animal experiments complied with Animal Research: Reporting of In Vivo Experiments (ARRIVE) guidelines and were carried out in accordance with the National Research Council’s Guide for the Care and Use of Laboratory Animals. The study was conducted with approval from the local Institutional Animal Care and Use Committee. *Bmal1*-stop-FL mice were generated on the C57BL/6J background as previously described [4; 5; 19]. Crosses of *Bmal1*-stop-FL mice with Alfp-Cre and Hsa-Cre lines generated mice with reconstitution of *Bmal1* in hepatocytes and cells that constitute skeletal muscle myo-fibers, respectively (LMRE). Experimental genotypes were: 1. wild type (WT) – *Bmal1^wt/wt^*, *Alfp-*cre^tg/0^, *Hsa-*cre^tg/0^; 2. *Bmal1* whole-body knockout (KO) – *Bmal1^stop-FL/stop-FL^*, *Alfp-*cre^0/0^, *Hsa-*cre^0/0^; 3. *Bmal1* liver and muscle reconstituted (LMRE) – *Bmal1^stop-FL/stop-FL^*; *Alfp*-cre^tg/0^; *Hsa*-cre^tg/0^. Additional experiments featured *Bmal1* liver-reconstituted mice (LRE) - *Bmal1^stop-FL/stop-FL^*; *Alfp*-cre^tg/0^. All experiments utilized male mice, age 8- to 14-week-old, entrained to a 12-h light: 12-h dark cycle. Mice were fed a standard chow diet with access either *ad libitum* (AL) or in a time-restricted manner only during the dark period from zeitgeber time (ZT) 12 to 24/0 (time-restricted feeding [TRF]).

### 2.2. Proteomic sample preparation

Preparation of liver and muscle for proteomic analysis was performed essentially as described previously [20; 21]. Briefly, tissue was lysed in 6 M guandine in 100 mM Tris, pH 8.5 containing 10 mM TCEP and 40 mM 2-choloroacetamide but tip-probe sonication and heated at 95℃ for 5 min. The samples were centrifuged at 20,000 x g for 30 min at 4℃ and supernatant containing protein was diluted with 1-volume of water and then precipitated with 4-volumes of acetone overnight at -30℃. Protein was pelleted by centrifugation at 4,000 x g for 5 min at 4℃ and washed with 80% acetone. The protein pellet was dried briefly to remove residual acetone and resuspended in 10% trifluoroethanol in 100mM HEPEs pH 7.5. Proteins were quantified with BCA, normalized to 10 µg/10 µL and digested overnight with 0.2 µg of sequencing-grade trypsin (Sigma) and 0.2 µg of sequencing-grade LysC (Wako, Japan) overnight at 37℃. Separate pooled samples of liver and muscle was also generated by mixing 4 µg of each sample and aliquoted into 10 µg/10 µL prior to digestion for use as internal controls. The peptides were labelled directly with 20 µg of 10-plex TMT (Thermo Scientific) for 1.5 h at room temperature. A total of six TMT labeling experiments (three for liver and three for muscle) were performed. The labelling scheme and experimental design is uploaded to the Pride Proteomics Repository (see Data Availability). After labelling, the reaction was deacylated with a final concentration of 0.2% hydroxylamine for 15 min followed by acidification to a final concentration of 1% trifluoroacetic (TFA) and the samples for each TMT experiment pooled. Samples were desalted directly with SDB-RPS microcolumns as previously described [21] and dried by vacuum centrifugation. Peptides were resuspended in 2% acetonitrile (MeCN) containing 0.1% TFA and fractionated by neutral pH reversed-phase chromatography on a Dionex 3500 micro-UHPLC. Peptides were injected onto a 0.3 mm x 15 cm column (C18BEH; 1.7 µm; Waters) maintained at room temperature and separated over 60 min gradient of 2-40% Buffer B at 6 µL/min into 48 fractions and automatically concatenated into 12 fractions in a looping fashion (Buffer A: 10 mM ammonium formate pH 7.5; Buffer B: 90% MeCN. Peptides were dried by vacuum centrifugation and stored at -80℃.

### 2.3. Proteomic analysis and data processing

Peptides were resuspended in 2% MeCN containing 0.1% TFA and separated on a nano-Dionex 3500 UHPLC. Peptides were injected onto a 0.075 mm x 40 cm column (C18AQ; 1.9 µm; Dr Maisch, Germany packed into PepSep, Marslev, Denmark)) maintained at 40 ℃ and separated over 60 min gradient of 2-30% Buffer B at 300 nL/min. Peptides were detected on an Orbitrap Eclipse mass spectrometer (ThermoFischer Scientific) via electrospray ionization in positive mode with 1.9 kV at 275 °C and RF set to 30%. The instrument was operated in data-dependent acquisition (DDA) mode with an MS1 spectrum acquired over the mass range 350–1,550 m/z (120,000 resolution, 8 x 10^5^ automatic gain control (AGC) and 50 ms maximum injection time) followed by MS/MS analysis with fixed cycle time of 3 s via HCD fragmentation mode and detection in the orbitrap (50,000 resolution; 1 × 10^5^ AGC, 86 ms maximum injection time and 0.7 m/z isolation width). Only ions with charge state 2-7 triggered MS/MS with peptide monoisotopic precursor selection and dynamic exclusion enabled for 30 s at 10 ppm. DDA Data were searched against the UniProt mouse database (October 2020; UP000005640_9606 and UP000005640_9606_additional) with MaxQuant v1.6.17.0 using default parameters with peptide spectral matches, peptide and protein false discovery rate (FDR) set to 1% [22]. All data were searched with oxidation of methionine set as the variable modification and carbamidomethylation of cysteine and 10-plex TMT of peptide N-termini and lysine set as fixed modifications. First search MS1 mass tolerance was set to 20 ppm followed by recalibration and main search MS1 tolerance set to 4.5 ppm, while MS/MS mass tolerance was set to 20 ppm. MaxQuant output data were initially processed with Perseus [23] to remove decoy data, potential contaminants and proteins only identified with a single peptide containing oxidized methionine.

### 2.4. Cell culture

Alpha mouse liver 12 (AML12) cells were purchased from ATCC (CRL-2254). Cells were cultured in Dulbecco’s modified Eagle’s medium: F12 (DMEM:F12, ATCC, 30-2006) supplemented with 10% fetal bovine serum (Corning, 35-015-CV), 1% penicillin/streptomycin, 1X insulin-transferrin-selenium (ITS) supplement (Corning, 25-800-CR), and 40 ng / mL dexamethasone. Cells were plated at confluency (40,000 cells per well) in Seahorse XF96 Cell Culture Microplates (Agilent). Cells were transfected with siRNA or mammalian expression vectors at the time of plating (reverse transfection). Cells were assayed 48 h later. AZD4547 (Abcam, ab216311) was administered 24 h prior to downstream measurements. The vehicle control (Veh) was 0.013% DMSO.

### 2.5. Oxygen consumption and extracellular acidification rates

Metabolic rates were measured using the Seahorse XFe96 Analyzer (Agilent) and Seahorse XF Cell Energy Phenotype Test Kit (Agilent, #103325-100) according to the manufacturer’s instructions.

### 2.6. FGF1 ELISA assay

Serum FGF1 protein was measured with the Mouse aFGF ELISA kit (RayBiotech, ELM- aFGF-1) according to the manufacturer’s instructions.

### 2.7. Western blot

Livers were homogenized in RIPA buffer (Thermo Scientific, J62524.AE) supplemented with protease inhibitors, lysed on ice for 10 min, sonicated (5 sec on, 5 sec off, for 4 cycles at 40% amp.), centrifuged at max speed at 4 °C for 10 min, and then the supernatant was collected. Twenty μg protein (or amount as otherwise indicated) was separated on a 4-20% gradient gel by SDS-PAGE and then transferred to a PVDF membrane. Blots were blocked for 1 hr with 5% milk in TBS with 0.1% tween-20 (TBST) at room temperature. Primary antibodies were diluted in 5% milk TBST and incubated with blots overnight at 4 °C (FGF1 – Abcam, ab207321; β-ACTIN – Abcam, ab8226; Albumin – Bethyl Laboratories, A90-134A). Blots were washed three times with TBST and then incubated with HRP-conjugated secondary antibodies (Millipore Sigma, AP160P and 12-348) for 1 h at room temperature. Blots were then washed three times, incubated with HRP substrate (Millipore, WBLUC0500) for 5 min at room temperature, and visualized using a chemiluminescent imaging system (Bio-Rad). Blots were quantified using ImageJ software. Where indicated, total protein on gels was visualized using stain-free gel technologies (Bio-Rad). For ex vivo secreted protein experiments, the concentrated supernatant entered the above protocol after the max speed centrifugation step.

### 2.8. Real-time quantitative PCR

RNA was extracted using the RNeasy Plus Mini Kit (Qiagen, 74136), and 500 ng of RNA was reverse transcribed using the Maxima First Strand cDNA Synthesis Kit for RT-qPCR (Thermo Scientific, K1641). qPCR was performed using PowerUP SYBR Green Master Mix (Applied Biosystems, A25742) on a QuantStudio 5 (Applied Biosystems). Gene expression data were normalized to 18S ribosomal RNA. Primer sequences were as follows: mouse Fgf1 forward 5’-ACACCGAAGGGCTTTTATACG-3’, reverse 5’-GTGTAAGTGTTATAATGGTTTTCTTCCA-3’.

### 2.9. Secretion of FGF1 from liver ex vivo

Secretion of proteins from liver was achieved similarly to previously published methods [24]. Mice were anesthetized with vaporized isoflurane (1-4%) and perfused (transcardiac) with 5 mL of PBS to remove blood from within the liver. The median lobe of the liver was harvested, washed in PBS, and transferred to a conical tube containing 5 mL of 1X CD CHO Medium (Gibco, 10743011) bubbled with carbogen. Liver was incubated for 1 h at 37 °C. The supernatant (the medium) was then concentrated using an Amicon Ultra-15 Centrifugal Filter Unit (Millipore Sigma, UFC901024) with a 10 KDa cutoff by swinging bucket centrifugation at 4,000 x g for 45 min at 4 °C. The resulting concentrated proteins were then prepared for western blot analysis.

### 2.10. Plasmids and siRNA

For knockdown experiments, cells were transfected with Silencer Select siRNAs (Thermo Fisher Scientific, IDs s65969 [Fgf1 #2], s65971 [Fgf1 #1], and Cat# 4390843 [negative control siRNA]) at a final concentration of 1 pmol using Lipofectamine RNAiMAX Transfection Reagent (Invitrogen, #13778150) according to the manufacturer’s instructions. For overexpression experiments, cells were transfected with 100 ng Fgf1 (NM_010197) Mouse Tagged ORF Clone (ORIGENE, MR201152) or control empty vector using Lipofectamine 3000 Transfection Reagent (Invitrogen, #L3000001) according to the manufacturer’s instructions.

### 2.11. Analysis of RNA-sequencing datasets

Analysis of mRNA was conducted using our previously published diurnal transcriptome datasets that were generated from the same cohort of mice as the proteomic data in this study or from a cohort of mice of identical age, sex, diet, and environmental conditions (GEO: GSE197726, GSE158600, GSE197455) [5; 12]. We used the differential rhythmicity and differential expression tool *dryR* [25] to identify rhythmic genes and genes with differential daily average expression between groups. *dryR* was performed on liver data from WT Alfp-Cre+ mice and *Bmal1* knockout mice and on muscle data from WT Hsa-Cre+ mice and *Bmal1* knockout mice. For the analysis, the starting set of genes oscillating in WT was defined as genes from rhythmic models 3, 4, and 5. Only genes with an mRNA expression value and a protein abundance value were analyzed.

### 2.12. Gene ontology analysis

Pathway enrichment analysis was carried out using the Database for Annotation, Visualization, and Integrated Discovery (DAVID) [26; 27]. Enrichments for biological process (BP), cellular compartment (CC), and molecular function (MF) were used where indicated. P<0.01 was considered statistically significant.

### 2.13. Statistical analyses

All data are displayed as mean ± S.E.M. unless otherwise noted. For each experiment, the number of biological replicates (n-value), statistical test and significance threshold can be found in the figure legend or main text. Complex statistical analyses of large-scale datasets are described within the corresponding methods section. Data were analyzed in Prism 6.0 software (GraphPad). The suitability of parametric vs non-parametric tests was determined by data distribution analysis tools in the software.

### 2.14. Data Availability

The proteomics data generated in this study are deposited to the ProteomeXchange Consortium (http://proteomecentral.proteomexchange.org/cgi/GetDataset) via PRIDE [28]

## 3. RESULTS AND DISCUSSION

### 3.1. Using tissue-specific reconstitution of Bmal1 to delineate layers of circadian control

To define the regulation of tissue proteomes by *Bmal1* and/or daily feeding rhythms, we used *Bmal1*-stopFL mice, which do not express the main transcriptional activator of the molecular clock, *Bmal1,* except in Cre recombinase-expressing cells [4; 19] (Figure 1A). *Bmal1*-stopFL mice lacking Cre (*Bmal1* knockout [KO]) are analogous to *Bmal1*-null mice and display severely impaired behavioral and molecular rhythms [4; 5; 19]. Hepatocyte-specific Alfp-Cre and skeletal muscle-specific Hsa-Cre lines were crossed to generate a single line in which *Bmal1* was reconstituted (i.e., rescued) in both hepatocytes and skeletal muscle (liver and muscle reconstituted [LMRE]) [12]. While this approach allowed us to analyze liver and muscle from the same mice, we considered that the abundance of some proteins may be influenced by *Bmal1* function in the other tissue, or by a synergistic effect of *Bmal1* in both tissues, rather than through rescue of local *Bmal1* function alone. We have previously shown that compared to wild-type (WT), KO and LMRE exhibit no significant changes in food intake, yet have a lower body weight and a higher fat-to-lean mass ratio [12].

**Figure 1.**
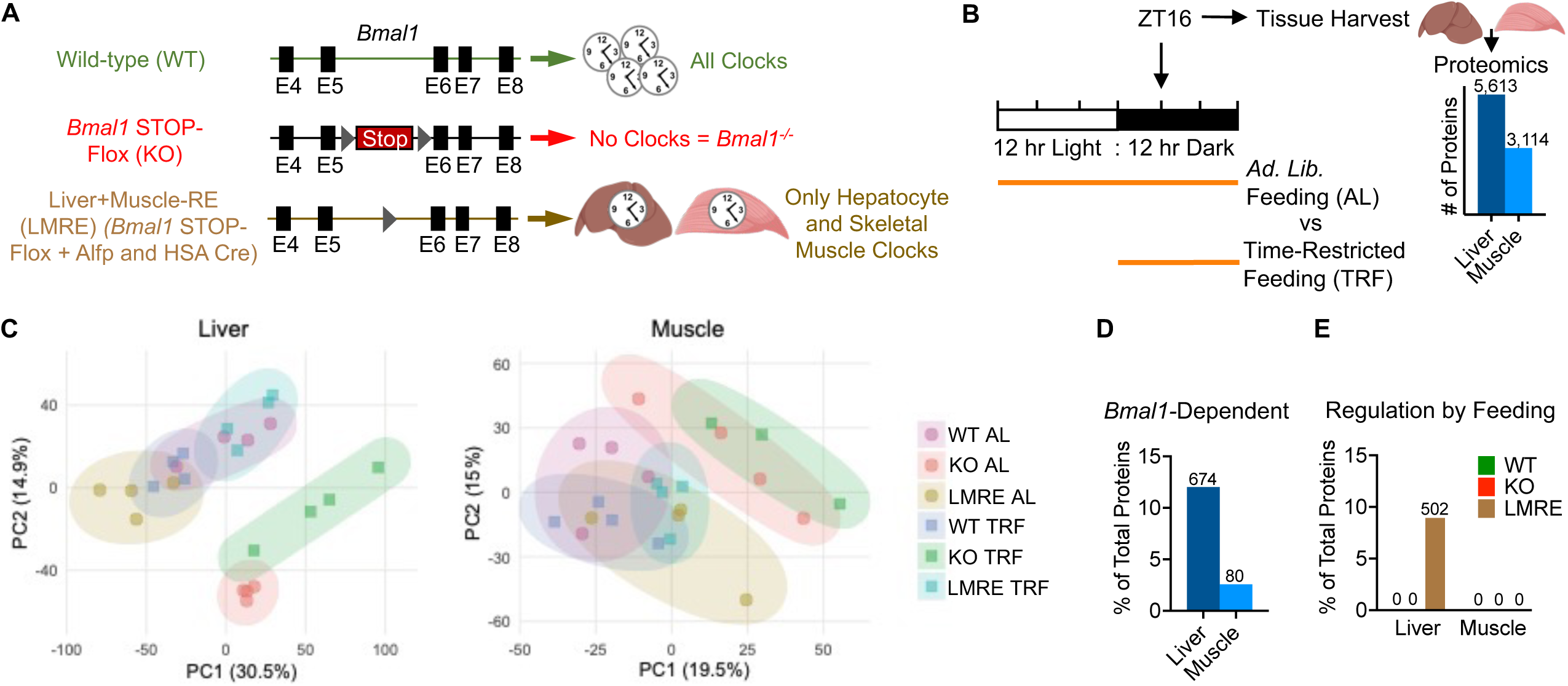
*Bmal1* remodels liver and skeletal muscle proteomes more potently than does time-restricted feeding. (A) Genetic scheme of cell-type-specific reconstitution of the molecular circadian clock. (B) Overview of experimental design. Proteomic analysis by Tandem Mass Tag (TMT) isobaric labelling combined with multi-dimensional liquid chromatography coupled to tandem mass spectrometry, n=4. Values on bars are number of proteins quantified. (C) Principal component analysis (PCA) of tissue proteomes. (D and E) Student’s t-test with false discovery rate correction, q<0.05, n=4. Values on bars are number of proteins. (D) Proteins up- or down-regulated in KO vs WT. (E) Proteins up- or down-regulated in TRF vs AL within each genotype. Figure 1A created with BioRender.com.

Due to lack of brain clock function [29], KO and LMRE mice lack robust 24-h rhythms of food intake under standard 12-h light/12-h dark conditions (Supplementary Figure S1A). To discern effects of physiological periods of feeding and fasting within and between genotypes, we placed mice under time-restricted feeding (TRF), with access to food only during the 12-h dark period (Figure 1B). Thus, we imposed a daily feeding-fasting behavior on KO and LMRE mice (Figure S1A), reinstating WT-like daily rhythms of energy expenditure and a daily switch between carbohydrate and lipid oxidation (respiratory exchange ratio), with no discernable differences between KO and LMRE [12]. We observed no significant changes in food intake in KO or LMRE mice under TRF, consistent with our previous reports [5; 12].

### 3.2. Bmal1 remodels the proteome more extensively than time-restricted feeding

Liver and gastrocnemius muscle tissues were harvested from WT, KO, and LMRE mice at zeitgeber time (ZT) 16 (4 h into the dark phase), from cohorts of both *ad libitum* (AL) or time-restricted feeding (TRF) mice, giving rise to six experimental groups for each tissue (n=4). This diurnal time point was identified as one of the peaks of rhythmic proteins in liver [16]. At this time, liver and muscle are engaged in the postprandial processing of macronutrients, when mice are normally active and feeding. Proteomic analysis was performed using Tandem Mass Tag (TMT) isobaric labelling combined with multi-dimensional liquid chromatography coupled to tandem mass spectrometry. Each tissue was analyzed with 3 x TMT 10-plex experiments that included a pooled reference across each experiment for normalization. This analysis quantified 5,613 proteins in liver and 3,114 proteins in muscle, with coverage across diverse cellular compartments (Figures 1B and S1B, Table S1).

To determine the global impact of *Bmal1* and feeding cycles on tissue proteomes, we performed principal component analysis (PCA). PCA plots revealed that liver proteomes segregated largely by the status of *Bmal1* and, to a much lesser extent, TRF (Figure 1C). This was also true for muscle, yet the effect of *Bmal1* KO was comparatively less than in liver. That is, the TRF used to reinstate normal feeding behavior did not appear to rescue the liver or muscle proteome based on global PCA. This contrasts with LMRE proteomes, which partially clustered towards WT.

We identified individual proteins affected by *Bmal1* by comparing WT to KO under AL conditions, and identified proteins regulated by feeding by comparing AL to TRF within each genotype (Student’s t-test, Benjamini-Hochberg corrected p-value [q-value] <0.05) (Table S1). The potency of *Bmal1* over feeding rhythms was also reflected in this analysis; whereas loss of *Bmal1* affected 12.01% of all liver proteins (Figure 1D), the maximum effect of feeding in any genotype was 8.94% (Figure 1E). In muscle, 2.57% of all proteins were affected by *Bmal1*, and remarkably, TRF failed to change the abundance of any proteins in muscle, even considering a less stringent q-value of <0.1. In liver, LMRE was the only genotype to respond substantially to TRF, with 8.94% of proteins affected. At q <0.1, we observed 113 (2.01%) and 7 (0.13%) proteins affected by TRF in KO and WT, respectively. This disparity between number of regulated proteins in liver and muscle remained when we also included proteins identified as *Bmal1*-dependent under TRF (Figures S1C and S1D). As feeding rhythms strongly affect gene expression in both WT and clock mutant mice [11; 15; 29; 30], it was intriguing that TRF had a minimal impact at the proteome level. Furthermore, the robust response to TRF in the livers of LMRE mice indicates that *Bmal1* substantially modulates the proteomic response to feeding.

### 3.3. Features of Bmal1-dependent proteomes in liver and skeletal muscle

To visualize *Bmal1*-dependent changes in more detail, we generated volcano plots of all proteins and highlighted top hits (Figure 2A). BMAL1, complexed with its binding partner CLOCK, is the main transcriptional activator of the molecular clock [31; 32], hence direct targets of BMAL1 are likely to be downregulated in its absence. Interestingly, we observed a nearly equal proportion of up- vs down-regulated proteins in KO liver and muscle compared to WT tissues. In line with the role of *Bmal1* in driving temporal metabolism [31], downregulated liver proteins were enriched for various lipid and carbohydrate pathways, and the top three downregulated muscle protein enrichments involved glucose (Figure 2B and Table S2). Of note, liver and muscle shared upregulated pathways related to circulatory homeostasis, such as blood coagulation and fibrinolysis (Figure 2B). Although most changes were indeed tissue-specific, we identified 22 *Bmal1*-dependent proteins common to liver and muscle (Figure 2C), including aldehyde dehydrogenase mitochondrial (ALDH2), an enzyme of alcohol metabolism that was reduced in KO and rescued in LMRE tissues, with no difference between AL and TRF (Figure 2D). Another interesting example was RAC-beta serine/threonine-protein kinase (AKT2), an enzyme of the insulin signal transduction pathway. Whereas AKT2 was upregulated in KO muscle, it was downregulated in KO liver, and rescued to WT levels in both LMRE tissues (Figure 2D). Such striking and disparate regulation likely stems from the tissue-specific outcomes of insulin signaling – insulin inhibits glucose production in liver and facilitates of glucose uptake into muscle [33; 34].

**Figure 2.**
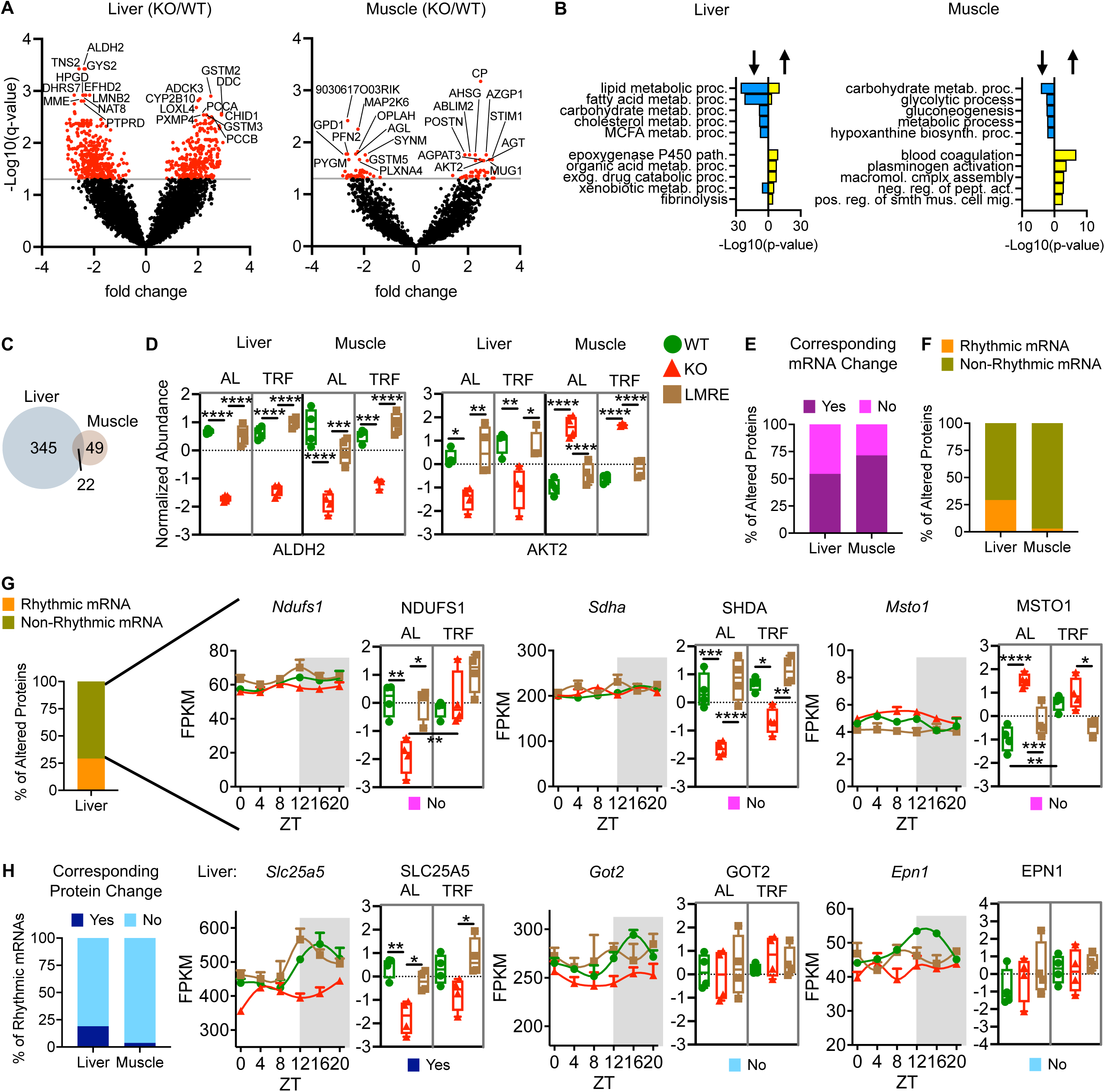
Features of *Bmal1*-dependent proteomes in liver and skeletal muscle. (A) Volcano plots highlighting *Bmal1*-dependent proteins, Student’s t-test with false discovery rate correction, q<0.05, n=4. Top 10 most significant up and down regulated proteins are indicated. (B) Gene ontology enrichment (biological process) analysis of proteins up-regulated (right protruding bars) and down-regulated (left protruding bars) in KO vs WT. (C) Venn diagram showing the overlap of *Bmal1*-dependent proteins in liver and muscle. Only proteins detected in both tissues were considered. (D) Examples of proteins regulated by *Bmal1* in both liver and muscle. AL – *ad libitum* feeding; TRF – time-restricted feeding; ALDH2 – aldehyde dehydrogenase 2, mitochondrial; AKT2 – RAC-beta serine/threonine-protein kinase. Two-way ANOVA with Tukey’s multiple comparisons test, * p<0.05, ** p<0.01, *** p<0.001, **** p<0.0001. (E-H) Comparisons of *Bmal1*-regulated proteins with their corresponding mRNA expression. WT and KO diurnal transcriptomes were analyzed with the differential rhythmicity algorithm *dryR*; rhythmic models were used to identify genes with rhythmic mRNA in WT and mean models (difference in daily average expression) were used to identify genes with altered mRNA levels between WT and KO. See also Table S3. Statistics on protein plots – two-way ANOVA with Tukey’s multiple comparisons test, * p<0.05, ** p<0.01, *** p<0.001, **** p<0.0001, n=4. (G) Examples of *Bmal1*-dependent liver proteins with non-rhythmic mRNA and no corresponding mRNA change in KO vs WT (i.e., post-transcriptionally regulated). NDUFS1 – NADH:ubiquinone oxidoreductase core subunit S1; SDHA – succinate dehydrogenase complex, subunit A, flavoprotein; MSTO1 – misato mitochondrial distribution and morphology regulator 1. Pink box refers to groupings from panel E. (H) Examples of liver genes with rhythmic mRNA peaking near ZT16 and their corresponding protein abundances at ZT16. Of note, the protein abundances of GOT2 and EPN1 break from their mRNA levels. SLC25A5 – solute carrier family 25, member 5; GOT2 – glutamatic-oxaloacetic transaminase 2, mitochondrial; EPN1 – epsin 1.

Through proteomics analysis, we identified the products of many recognizable clock-controlled genes whose transcriptional rhythms are well documented as well as novel targets whose regulation may or may not originate from changes in transcription. Many regulatory steps separate nascent transcription from steady-state protein abundances (e.g., mRNA half-life, RNA processing events, translation efficiency). To tease apart whether the observed protein changes stemmed from changes in mRNA, we used *dryR* [25] mean models to identify genes whose average daily mRNA levels were differentially regulated between WT and *Bmal1* KO mice (Table S3). For this analysis, we used transcriptome data from our previous study that featured identical experimental conditions yet with transcriptomic data obtained at six timepoints over the light-dark cycle [5; 12]. Rather than limit tests to ZT16, this approach of comparing average daily RNA levels takes into consideration that protein abundances do not necessarily go lockstep in time with RNA and may be slightly delayed due to the added timeframe of translation. The analysis revealed that 54.34% of affected liver proteins and 71.43% of affected muscle proteins had a corresponding change in average mRNA levels (Figure 2E). These data implicate that posttranscriptional regulation has a larger role in shaping protein abundances in liver than muscle.

Next, we queried which *Bmal1*-regulated proteins at ZT16 were the product of rhythmic genes, which we defined using *dryR* rhythmic models of the WT transcriptomes of both tissues (Table S3). Surprisingly, we found that most affected proteins were the product of non-rhythmic genes (70.77% in liver; 97.14% in muscle; Figure 2F). Focusing on liver, we found that interrogation of functional enrichments unique to each group of proteins revealed that affected proteins of non-rhythmic genes were associated with mitochondrion organization and morphogenesis, among other pathways (Figures 2G and S2A). Notable examples were DNA polymerase subunit gamma-2 mitochondrial (POLG2), which promotes mitochondrial DNA synthesis, mitofusin-2 (MFN2), a GTPase essential for mitochondrial fusion, and many subunits of ubiquinone NADH dehydrogenase (NDUF, complex I), adding mechanistic insight to previous studies reporting a role for *Bmal1* in daily mitochondrial homeostasis [35].

Conversely, we identified genes with rhythmic mRNA and asked whether the abundance of their corresponding protein was altered by *Bmal1* knockout. We included genes with peak expression within the dark phase (ZT14 to ZT22), the time when protein measurements were made. We found that only 19.01% of rhythmic liver genes and 3.70% of rhythmic muscle genes had proteins that were altered with loss of *Bmal1* (Figure 2H). Unaffected proteins either represent genes whose rhythmicity is lost at the protein level or genes whose protein abundance lags behind mRNA abundance to a large extent. In liver, this group of proteins was uniquely enriched for gene ontology terms related to translation (Figure S2B). In contrast, rhythmic genes that indeed had proteins that were altered by loss of *Bmal1* were uniquely enriched for pathways of energy metabolism. Together, these data reveal that alterations in mRNA abundance upon loss of BMAL1 are not reflected in the protein abundance, and *vice versa*, underscoring a complex regulation of these aspects of cellular control by BMAL1.

### 3.4. Partial rescue of the liver ensemble proteome by hepatocyte Bmal1

Next, we defined restoration of the liver proteome in LMRE mice. In cases where proteins were not rescued, we provide two explanations that inform on the organization the clock network at the systemic and local levels: 1) hepatocyte *Bmal1* is insufficient to maintain the abundance of that protein and extrinsic signals tied to *Bmal1* function elsewhere are needed, or 2) the protein is primarily controlled by *Bmal1* in a different cell type. Considering hepatocytes constitute the bulk of liver cells as well as liver protein [36], and that the average daily expression of *Bmal1* in LMRE liver is similar to WT liver [4; 5] (i.e., recombination efficiency is high and hepatocytes produce the bulk of *Bmal1* transcripts in liver), we expect explanation one to apply in most cases. However, non-parenchymal cells such as endothelial cells, stellate cells and Kupffer cells contribute to the liver proteome [36; 37], thus explanation 2 also has merit. As many functions are unique to hepatocytes, a functional analysis of proteins helped us to further tease apart the contribution of hepatocytes. PCA plots showed that AL proteomes of WT and LMRE livers were similar but did not overlap completely (Figure 1C).

To identify the proteins responsible for this divergence, we performed individual statistical comparisons on each protein and split *Bmal1*-dependent proteins into categories of rescued (q<0.05, WT vs KO, LMRE vs KO; q>0.05, WT vs LMRE) and non-rescued (q<0.05, WT vs KO; q>0.05, LMRE vs KO, WT vs LMRE) (Figures 3A, 3B, and S3A). We observed a near equal proportion of rescued (317) vs non-rescued (357) proteins, for both up and downregulated proteins. In WT and LMRE under TRF, non-rescued proteins appear more similar (Figure S3A). However, the effect of TRF was seemingly equally due to a reduction of protein abundances in WT and an increase in protein abundances in LMRE. Although TRF rescued certain proteins in KO tissues, abundances remained dysregulated overall (Figure S3A).

**Figure 3.**
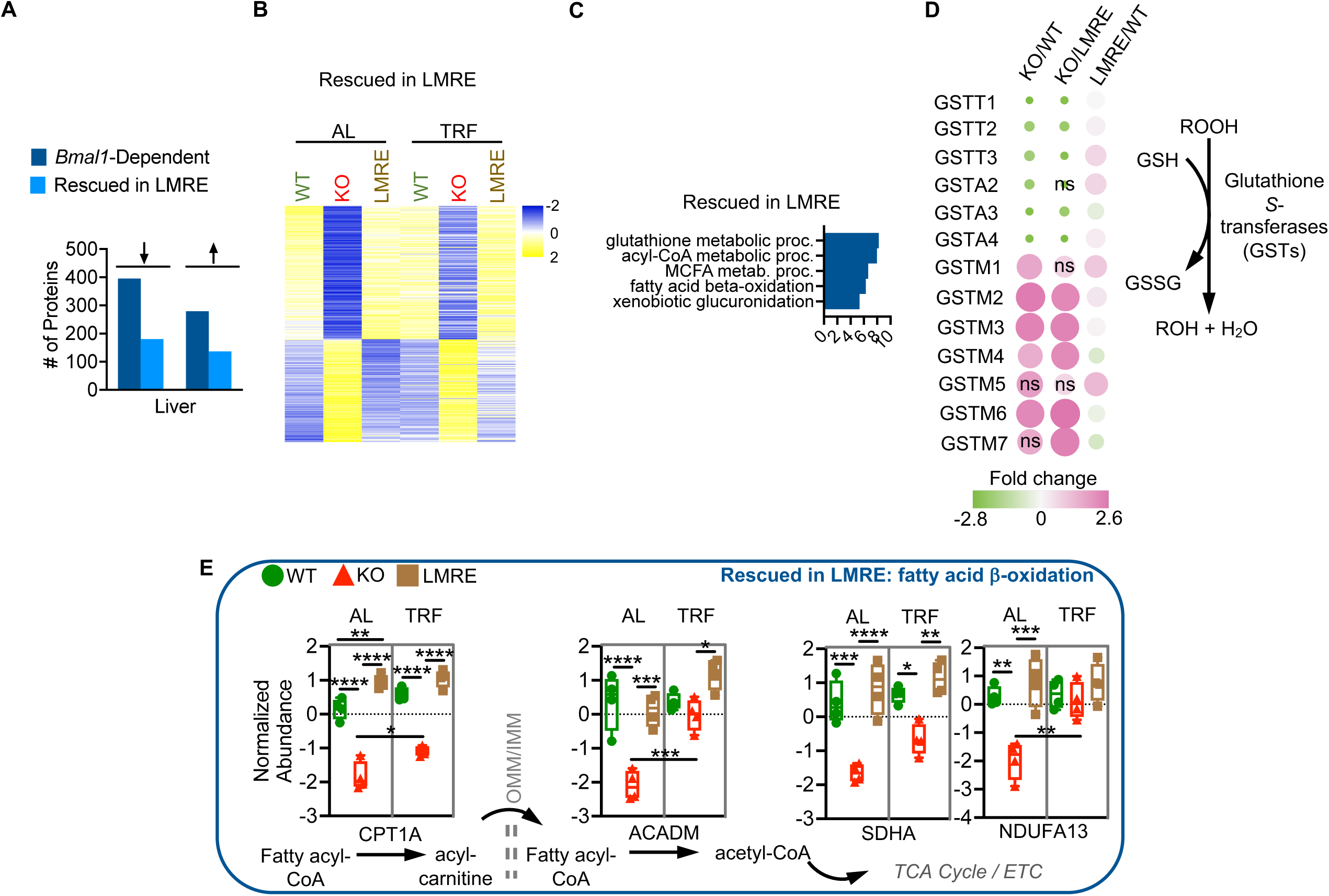
Partial rescue of the liver ensemble proteome by hepatocyte *Bmal1*. (A-E) *Bmal1*-dependent proteins (WT vs KO, q<0.05) were statistically categorized by false-discovery-rate-corrected p-values (q-values) from Student’s t-tests. Rescued in LMRE = WT vs KO, q<0.05; LMRE vs KO q<0.05; WT vs LMRE q>0.05. Arrows indicate up- or down-regulation in KO vs WT. (B) Heatmaps of rescued proteins. (C) Gene ontology enrichment (biological process) analysis showing the top 5 uniquely enriched pathways for each category of proteins. See also Table S2. (D) Bubble plot of quantified glutathione *S*-transferases (GSTs) of the theta (T), alpha (A), and mu (M) classes. Bubble color and size are proportional to fold change, ns – not significant. ROOH – hydroperoxide functional group. GSH – reduced glutathione; GSSG – oxidized glutathione; (E) Simplified scheme showing key steps of fatty acid β-oxidation. Two-way ANOVA with Tukey’s multiple comparisons test, * p<0.05, ** p<0.01, *** p<0.001, **** p<0.0001, n=4. OMM/IMM – outer mitochondrial membrane/inner mitochondrial membrane; ETC – electron transport chain; MUFAs – monounsaturated fatty acids; CPT1A – carnitine palmitoyltransferase 1a, liver; ACADM – acyl-coenzyme A dehydrogenase, medium chain; SDHA – succinate dehydrogenase complex, subunit A, flavoprotein; NDUFA13 – NADH:ubiquinone oxidoreductase subunit A13.

Specific functions of rescued proteins were revealed by mapping proteins to pathways (Figure 3C). Consistent with our previous studies [4; 5], rescued proteins were involved in functions such as NAD^+^ metabolism, involving the salvage pathway enzymes NAMPT and NMNAT3, and glycogen metabolism through GYS2, which are highly rhythmic processes in mouse liver. In line with the reasoning that non-rescued proteins are regulated by extrinsic signals, “response to nutrient” was among the top 5 enrichments of this category (Figure S3B). We identified proteins involved in responding to xenobiotics (GCLC), energetic state (PKLR, GATM) specific energy substrates (HMGCL, AACS) and signaling activities (PTEN, ADIPOQ). Interestingly, top hits included enzymes involved in branched-chain amino acid catabolism (ACAD, HIBADH, IVD, BCKDHB), a function intertwined with skeletal muscle [38] (Figure S3C). Glucose transporter 2 (GLUT2) was an example of partial rescue, showing a significant up regulation in LMRE liver versus KO under both *ad libitum* and TRF conditions, yet not reaching WT abundance (Figure S3C).

Several pathways were remarkably distinct in terms of rescue and highlighted intricacies of daily liver metabolism. Glutathione is a major antioxidant that detoxifies reactive oxygen species, xenobiotics, and peroxides. Whereas GCLC, the rate-limiting enzyme for glutathione synthesis, was not rescued in LMRE (Figure S3C), nearly all glutathione *S*-transferases (GSTs) were (Figure 3D). GSTs use glutathione to reduce, and in effect detoxify, diverse substrates [39]. We observed full rescue of the abundance of GST proteins belonging to the alpha, mu, and theta classes. Intriguingly, only mu proteins were upregulated in *Bmal1* KO livers (Figure 3D), indicating dichotomy in the directionality of regulation among GSTs, which may relate to substrate specificity. Glutathione rhythms are reported to anticipate their need, peaking just prior to the intake of potentially harmful substances during feeding at the onset nighttime [40; 41]. Our data showed that GST abundances were unaffected by TRF (Figure S3D). Together, these data are indicative of extrinsic control of the machinery for glutathione production and intrinsic clock control of GSTs, helping to explain the roles of the liver clock and feeding rhythms in controlling chronotoxicity in the liver [42; 43].

“Lipid metabolic process” was the top enrichment for both rescued and non-rescued proteins (Table S2), but a more detailed analysis of this large pathway revealed that fatty acid oxidation was enriched in rescued proteins and the opposing process of fatty acid synthesis was enriched in non-rescued proteins (Figures 3C, S3B, and S3E). The liver is a major hub in organismal lipid metabolism. Fatty acids are oxidized as energy substrates, synthesized from carbohydrate building blocks, stored, or circulated to other tissues in the form of triglycerides. These activities exhibit circadian rhythms to anticipate opportunities to store energy during feeding (*de novo* lipogenesis) and utilize energy stores during periods of fasting (fatty acid β-oxidation) [6; 44; 45]. Enzymes mediating key steps in peroxisomal and mitochondrial fatty acid oxidation were rescued in LMRE livers (Figures 3C and 3E), including carnitine palmitoyltransferase 1 a (CPT1A), which catalyzes the rate-limiting step for long-chain fatty acids, and acyl-coenzyme a dehydrogenase medium chain (ACADM), which acts on medium-chain fatty acids. Mitochondrial proteins that support energy production downstream of β-oxidation were also numerous among fully rescued proteins. In contrast, enzymes mediating key steps in *de novo* lipogenesis were classified as non-rescued (Figure S3B and S3E). The rate-limiting step in *de novo* lipogenesis is the carboxylation of acetyl-CoA to malonyl-CoA by acetyl-CoA carboxylase (ACC). ACC alpha (ACACA) abundance was similarly downregulated in KO and LMRE livers under AL conditions. This was also true for ATP citrate synthase (ACLY), which produces acetyl-CoA upstream of ACC, and for fatty acid synthase (FASN), which synthesizes palmitate downstream of ACC. These changes were concomitant with lower abundance of sterol regulatory element-binding protein 1 (SREBF1, also called SREBP), a transcription factor that promotes lipogenesis [46] (Figure S3F). As these data were acquired at ZT16, a time point characterized by higher *de novo* lipogenesis activity [6; 46], the protein abundances demonstrate that extrinsic signals support fatty acid-synthesizing machinery. Our data show that one potential extrinsic signal – feeding – tended to normalize protein abundances among the genotypes, but this effect was partially driven by a reduction in protein abundances in WT. These results highlight the importance of non-autonomous regulation by the clock system on hepatic lipogenesis.

### 3.5. Partial rescue of the muscle ensemble proteome by myofiber Bmal1

Next, we defined the ability of *Bmal1* in muscle fibers to rescue the skeletal muscle proteome. Recent single-nucleus sequencing studies show that ∼70% of nuclei within skeletal muscle are the muscle-fiber associated myonuclei [47]. Here, we used Hsa-Cre to rescue *Bmal1* specifically in such myonuclei. Muscle fibers are the sites of metabolic activity in the muscle, and we have previously shown that *Hsa*-Cre recombination is highly efficient in LMRE muscles [12]. Accordingly, our model is suitable to investigate proteomic responses of these cells. Like liver, non-rescued proteins in muscle may be produced by non-myolineage cell types or regulated by missing extrinsic signals from other tissue clocks.

We statistically categorized muscle proteins as rescued or non-rescued and visualized the results with heatmaps (Figures 4A, 4B, and S4A). Fifteen of the 18 proteins rescued in LMRE were downregulated in KO compared to WT. This directionality was not observed in liver or for non-rescued proteins in muscle. Due to the small number of proteins, we did not observe any significant functional enrichments for rescued proteins (Figure 4C). Notable proteins included glutamic pyruvic transaminase (GPT, also known as alanine aminotransferase 1 [ALT]), which serves a key function in muscle carbohydrate metabolism by converting glycolysis-derived pyruvate to alanine, 5-oxoprolinase (OPLAH), an ATP-hydrolyzing enzyme involved in glutamate production, and mitogen-activated protein kinase kinase 6 (MAP2K6), a regulator of MAP kinase (Figure 4C and 4D).

**Figure 4.**
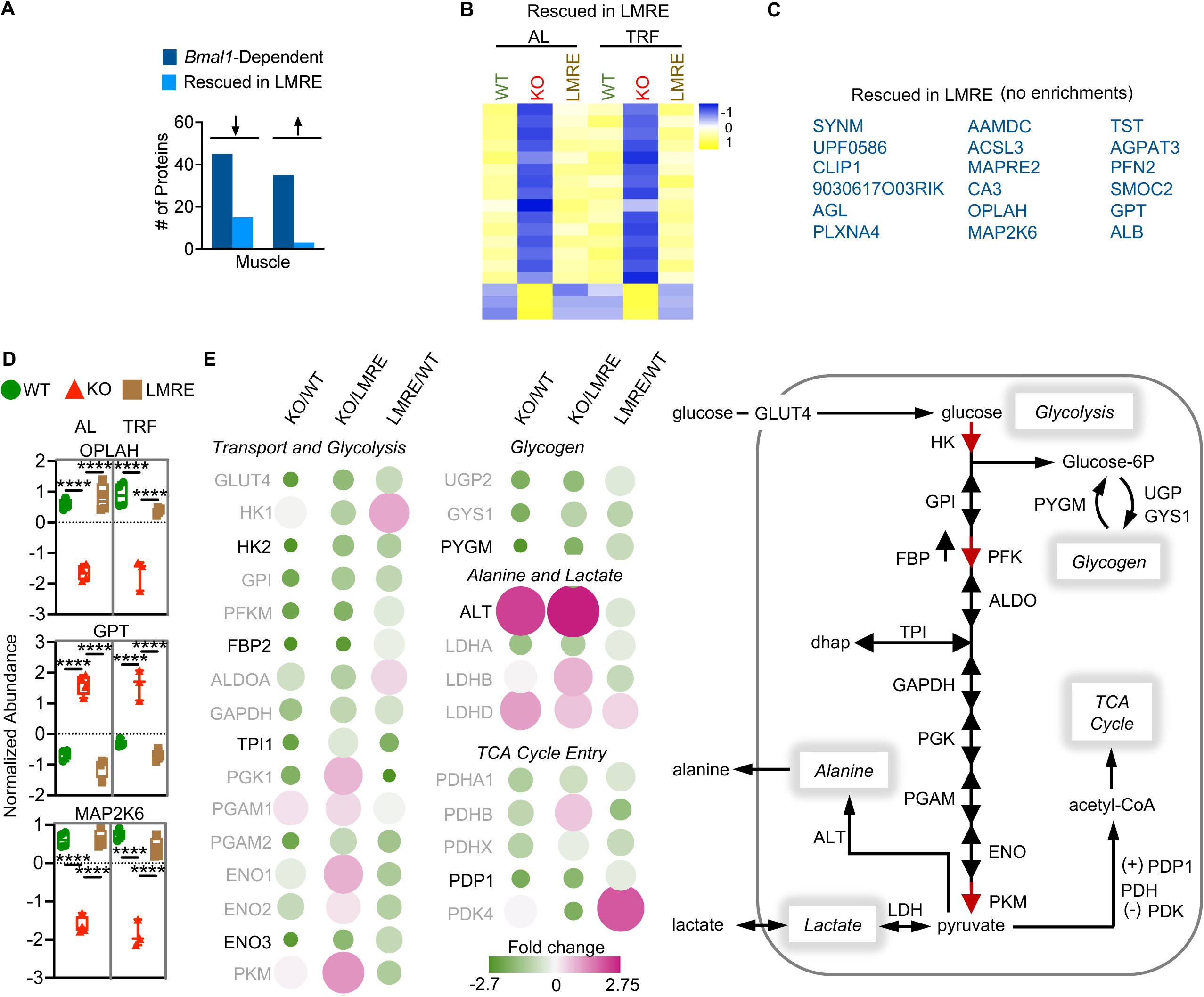
Partial rescue of the skeletal muscle ensemble proteome by myofiber *Bmal1*. (A-F) *Bmal1*-dependent proteins (WT vs KO, q<0.05) were statistically categorized by false-discovery-rate-corrected p-values (q-values) from Student’s t-tests. Rescued in LMRE = WT vs KO, q<0.05; LMRE vs KO q<0.05; WT vs LMRE q>0.05. Arrows indicate up- or down-regulation in KO vs WT. (B) Heatmaps of rescued proteins. (C) Gene ontology enrichment (biological process) analysis showing the top 5 uniquely enriched pathways for each category of proteins. See also Table S2. (D) Examples of rescued proteins in muscle of LMRE mice. Two-way ANOVA with Tukey’s multiple comparisons test, **** p<0.0001, n=3-4. OPLAH – 5-oxoprolinase; GPT (ALT) – alanine aminotransferase 1; MAP2K6 – mitogen-activated protein kinase kinase 6. (E) Left, bubble plot of quantified proteins that carry out key steps in glucose metabolism. Bubble color and size are proportional to fold change. Non-significant changes have a grayed-out protein name. Right, simplified scheme showing key steps in muscle glucose metabolism. GLUT4 (SLC2A4) – solute carrier family 2 member 4; HK – hexokinase; GPI – glucose-6-phosphate isomerase; PFKM – phosphofructokinase, muscle; FBP2 – fructose bisphosphate 2; ALDOA – aldolase A; GAPDH – glyceraldehyde-3-phosphate dehydrogenase; TPI1 – triosephosphate isomerase 1; PGK1 – phosphoglycerate kinase 1; PGAM – phosphoglycerate mutase 1; ENO – enolase; PKM – pyruvate kinase, muscle; UGP2 – UDP-glucose pyrophosphorylase 2; GYS1 – glycogen synthase 1; PYGM – muscle glycogen phosphorylase; ALT (GPT) – alanine aminotransferase 1; LDH – lactate dehydrogenase; PDH – pyruvate dehydrogenase; PDP1 – puruvate dehydrogenase phosphatase catalytic subunit 1; PDK4 – pyruvate dehydrogenase kinase, isoenzyme 4; 6P – 6-phosphate; dhap – dihydroxyacetone phosphate.

Two of the top five functional enrichments of non-rescued proteins involved glucose metabolism (Figure S4B). During feeding, most glucose is cleared from the blood via insulin-induced uptake into skeletal muscle [48; 49]. This process is under circadian control; the central clock in the brain drives mice to consume most food during the dark active phase [50], BMAL1 modulates glucose-stimulated insulin release from pancreatic beta cells [8], and muscle BMAL1 times the expression of key enzymes to meet the demand [51]. By tracing ^13^C-labeled glucose in LMRE mice, we have recently shown that although muscle BMAL1 is necessary for maximal glucose uptake and oxidation, it is not sufficient [12], and the addition of a liver clock in *ad libitum* conditions does not augment this function. In line with this finding, we observed that protein abundances for several key glucose enzymes were still deregulated in LMRE mice (Figure 4E).

Glucose has several fates in muscle: 1) oxidation via the TCA cycle, 2) conversion to lactate or alanine as a glycolytic endpoint, 3) storage as glycogen, and 4) release as succinate [48; 52-54] (Figure 4E). Oxidation supports energy production, glycogen supplies glycolytic intermediates (and substrates for energy production), and lactate and alanine provide the liver with substrates for the TCA cycle, gluconeogenesis, and the disposal of nitrogen. Several glycolysis enzymes, including the rate limiting enzyme hexokinase 2 (HK2), showed a trend for partial restoration in LMRE muscles (Figure 4E). The muscle isoform of the primary glycogenolysis enzyme, glycogen phosphorylase muscle associated (PYGM), also appeared to be partially rescued, along with pyruvate dehydrogenase phosphatase catalytic subunit 1 (PDP1) (Figure 4E). PDP1 dephosphorylates and activates the pyruvate dehydrogenase complex, generating acetyl-CoA that feeds into the TCA cycle. These data suggest that the abundances of enzymes responsible for processing glucose and directing the flow of its carbons are more strongly supported by muscle-extrinsic functions of *Bmal1*. Because their abundances still showed tendencies for improvement as compared to KO muscle, we posit that the muscle-extrinsic functions of BMAL1 fine-tune or provide robustness to these pathways.

### 3.6. An interaction between Bmal1 and feeding dictates ribosomal protein abundances in liver

TRF paradigms substantially modulate mRNAs, especially in clock mutant mice that lack a typical feeding-fasting rhythm under *ad libitum* conditions [11; 15; 29; 30]; however, changes in protein abundances in response to TRF are less-well characterized. When comparing AL vs TRF within genotypes, we observed that LMRE was the only genotype to respond robustly to TRF (502 proteins, q<0.05), and this only occurred in liver (Figure 5A). There were no proteins affected in WT or KO at this significance threshold. Trends were maintained if the threshold was eased to q<0.1, with muscle still lacking any feeding-responsive proteins, irrespective of genotype. This global analysis indicates that at the protein level, the liver, but not the muscle, is a major target of TRF.

**Figure 5.**
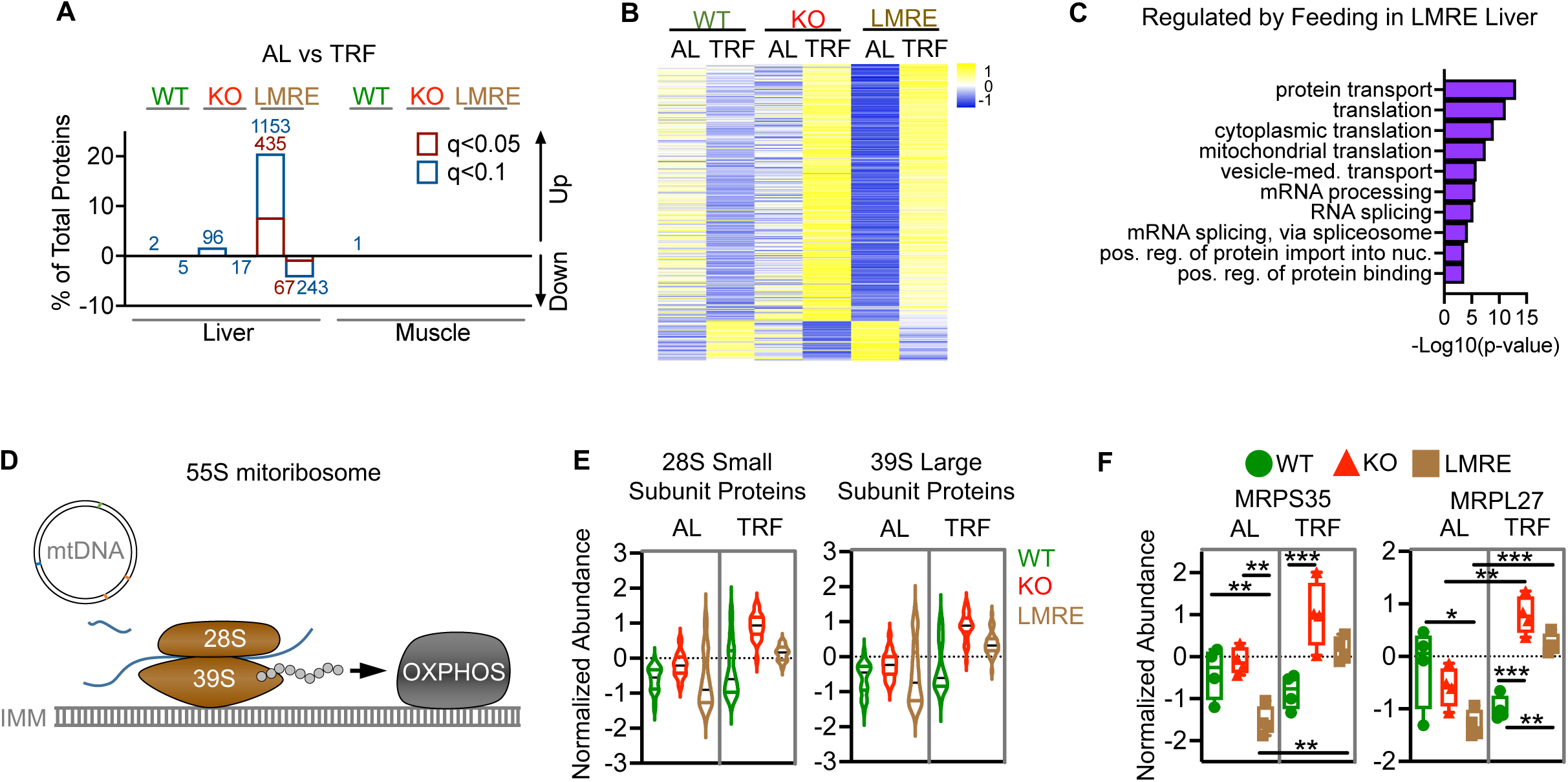
An interaction between *Bmal1* and feeding rhythms dictates ribosomal protein abundances in liver. (A) Effect of feeding on protein abundances within genotypes. AL – *ad libitum* feeding; TRF – time-restricted feeding. Student’s t-test with false discovery rate correction, q<0.05, n=4. Results using a less-stringent q-value (0.1) are shown for comparison. Values on bars are number of proteins. (B-C) Analysis of liver proteins affected by TRF (q<0.05) in LMRE. (B) Heatmap. (C) Gene ontology enrichment (biological process) analysis showing the top 10 enriched pathways. (D) simplified scheme of the mitoribosome and its function in the mitochondrion. The small (28S) and large (39S) subunits form the 55S mitoribosome that translates RNA produced from mitochondrial DNA (mtDNA). RNAs include those that encode proteins of the complexes of the electron transport chain, which enables oxidative phosphorylation (OXPHOS). IMM – inner mitochondrial membrane. (E) Violin plot showing the average normalized abundance of 27 small subunits (28S, left) and 45 large subunits (39S, right) in liver. (F) Example abundances of small and large mitoribosome subunits in liver. Two-way ANOVA with Tukey’s multiple comparisons test, * p<0.05, ** p<0.01, *** p<0.001, n=4. MRPS35 – mitochondrial ribosomal protein S35; MRPL27 – mitochondrial ribosomal protein L27.

A heatmap of the 502 feeding-responsive proteins from LMRE livers revealed interesting patterns of protein abundance between the genotypes (Figure 5B). In KO and LMRE, most proteins were upregulated by TRF, but changes were only significant in LMRE because changes in KO were considerably smaller in magnitude. Conversely, the same proteins were modesty and non-significantly downregulated by TRF in WT. WT mice already enact a robust feeding-fasting rhythm, and we reveal that a slight enhancement of this behavior by TRF had only a minimal impact, even if thousands of genes at the transcript level are affected by the same TRF paradigm in WT mice [5]. Similarly, TRF can positively impact the transcriptome of *Bmal1* KO livers [15; 29]. However, these current data indicate quite different protein level responses to TRF during feeding, suggesting a buffering capacity, particularly in co-expressed genes [55]. From these results, we concluded that liver *Bmal1* and extra-hepatic *Bmal1* augment the liver’s response to TRF.

We identified three main cellular activities that dominated the feeding-responsive proteins in liver: protein transport, translation, and RNA processing (Figure 5C). Previous studies have elucidated mechanisms by which translation is controlled by the clock and feeding [13; 14; 56]. In mouse livers, a subset of mRNAs displays increased translation during the dark phase, facilitated by increased ribosome biogenesis that is aligned with feeding time [14]. On one hand, the molecular clock shapes this rhythm by controlling transcription of translation initiation factors, ribosomal proteins and ribosomal RNAs [56]. On the other hand, daily feeding rhythms drive nutrient-responsive signaling pathways, and nutrient status, in turn, is a potent regulator of translation [13]. Here, considering translation machinery and regulatory proteins under *ad libitum* conditions, we did not observe any enriched functions related to translation in *Bmal1* knockout livers (Table S2). Instead, enrichment of translation gene ontology terms was specific to conditions in which feeding-fasting cycles were engaged. A more detailed analysis revealed that only one ribosomal protein (RPL13) was affected by *Bmal1* KO (under *ad libitum* feeding) whereas 62 were regulated by feeding (Figure S5A). Proteins constituting both the large (60S) and small (40S) ribosomal subunits were affected. Several members of the eukaryotic initiation factor (EIF) protein family were upregulated by TRF in LMRE. These increased protein abundances may indicate increased translation, and our analysis was performed in a diurnal phase associated with maximal translation, ribosome accumulation, and typical feeding times. Therefore, our data suggest that circadian control of feeding behavior is an important driver of translation in liver.

Analysis of feeding-responsive proteins also illuminated a previously unappreciated effect on mitochondrial translation. In addition to cytoplasmic ribosomes, a distinct set of ribosomal proteins constitutes two ribosome subunits assembled within the mitochondria [57] (Figure 5D). One 28S small subunit and one 39S large subunit comprise a mitochondrial ribosome (the 55S mitoribosome). We found that average expression levels of 28S and 39S mitochondrial ribosomal proteins exhibit a similar pattern of abundance as the individual proteins we identified as statistical hits (Figure 5E and 5F). The constituent ribosomal proteins are encoded by nuclear DNA and support translation of proteins encoded by the mitochondrial DNA. This process is critical for proper oxidative phosphorylation, as the protein coding genes of mitochondrial DNA produce subunits of complex I, IV, and V of the electron transport chain [58]. The fact that these changes were only observed in LMRE suggests that local clocks are required to transduce signals from feeding to regulate mitochondrial translation.

### 3.7. Secreted proteins are under BMAL1 control in liver and muscle

Liver and muscle cells also produce extracellular or secreted proteins. Secreted proteins modulate a wide range of functions through autocrine, paracrine, and endocrine mechanisms. In liver, it has been shown that secreted proteins accumulate during nighttime and are released in the middle of the day [16]. The timing of our proteomics analysis corresponded with the known time of accumulation of secreted proteins within the liver (ZT16, nighttime) [16], therefore we used the opportunity to identify secreted proteins regulated by *Bmal1* and TRF.

Most proteins are secreted via the classical/conventional secretory pathway – synthesized by endoplasmic reticulum (ER)-tethered ribosomes, transferred in vesicles to the Golgi apparatus, packaged into secretory vesicles, and trafficked to the plasma membrane for release via exocytosis [59; 60]. Proteins downregulated in *Bmal1* KO livers were enriched for ER to Golgi vesicle-mediated transport, lipoprotein transport, and vesicle mediated transport (Table S2), suggesting that BMAL1 supports protein secretion. Several components of coat protein complex II (COPII), which facilitates the formation of vesicles for ER to Golgi transport, were down-regulated in *Bmal1* KO and rescued in LMRE (including secretion associated ras related GTPase 1b [SAR1B]; sec16 and 31 homologues a [SEC16A, SEC31A]) (Figure 6A), as were components of coat protein complex I (COPI), which mediates retrograde Golgi to ER transport. These changes were not observed in muscle (Figure 6A).

**Figure 6.**
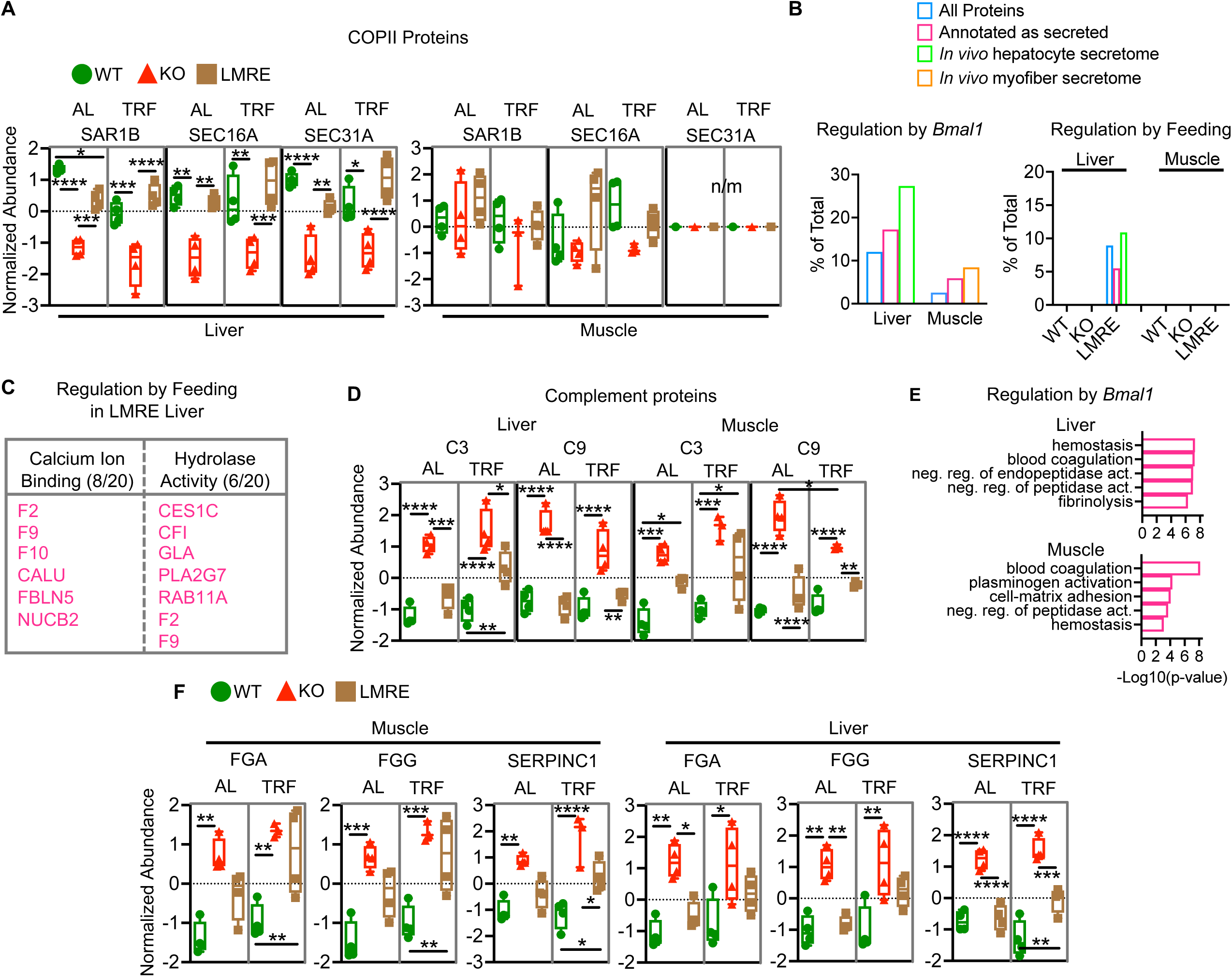
Secreted proteins are targeted by *Bmal1* in liver and skeletal muscle. (A) Example proteins from the coat protein two (COPII) complex, a mechanism of secreted proteins which enables the formation of vesicles to transport proteins from the endoplasmic reticulum to the golgi apparatus. Two-way ANOVA with Tukey’s multiple comparisons test, * p<0.05, ** p<0.01, *** p<0.001, **** p<0.0001, n=4 (n=3 for TRF KO muscle). AL – *ad libitum* feeding; TRF – time-restricted feeding. n/m – not measured; SAR1B – secretion associated Ras related GTPase 1B; SEC16A – SEC16 homolog A, endoplasmic reticulum export factor; SEC31A – Sec31 homolog A. (B-F) Analysis of secreted proteins. (B) Student’s t-test with false discovery rate correction, q<0.05, n=4, WT vs KO (regulation by *Bmal1*), AL vs TRF within genotype (regulation by feeding). Displayed as percentage of detected proteins in that class. Secreted proteins either annotated as secreted via Uniprot or published *in vivo* secretomes for hepatocytes or myofibers (Wei et al., 2023). (C) Gene ontology enrichment (molecular function) analysis of secreted proteins regulated by feeding in LMRE liver. Numbers indicate hits/total proteins in class. F – coagulation factor; CALU – calumenin; FBLN5 – fibulin 5; NUCB2 – nucleobindin 2; CES1C – carboxylesterase 1C; CFI – complement component factor i; GLA – galactosidase, alpha; PLA2G7 – phospholipase A2; RAB11A – RAB11A, member RAS oncogene family. (D) Analysis of complement proteins in liver and muscle. C3 –Complement component 3; C9 – Complement component 9. Two-way ANOVA with Tukey’s multiple comparisons test, * p<0.05, ** p<0.01, *** p<0.001, **** p<0.0001, n=4. (E) Gene ontology enrichment analysis showing the top 5 enriched pathways. (F) Analysis of coagulation cascade proteins in liver and muscle. FGA/G – fibrinogen alpha and gamma chains, SERPINC1–serine protease inhibitor c 1. Two-way ANOVA with Tukey’s multiple comparisons test, * p<0.05, ** p<0.01, *** p<0.001, **** p<0.0001, n=4.

To interrogate secreted proteins from our dataset, we either filtered for “extracellular” gene ontology classification or used published *in vivo* secretomes for hepatocytes and myofibers [61]. In both cases, we found secreted proteins were more heavily affected by *Bmal1* KO than the total protein pool, suggesting an importance of this group of proteins among clock-controlled genes (Figure 6B). LMRE liver was the only group to exhibit changes in secreted proteins in response to TRF (Figure 6B). Of the 30 feeding-responsive proteins annotated as secreted in LMRE, 10 were also modulated by BMAL1. Among the 20 feeding-responsive proteins were many zymogens and proenzymes, inactive precursor peptides that require further cleavage to become active (Figure 6C). Zymogens are key steps in blood coagulation and complement (immune system) cascades [62]. The thrombin precursor prothrombin (F2), the plasmin precursor plasminogen (PLG), the kinin precursor kininogen (KNG2), and the coagulation factors IX and X (F9 and F10) were upregulated by TRF in LMRE livers. The same trend was observed in KO livers, but increases were not statistically significant, implying that *Bmal1* is required to enhance the responses of these proteins to feeding.

Notably, complement cascade proteins were strongly upregulated in both liver and skeletal muscle of *Bmal1* KO mice (Figures 6D). Of the 23 detected complement proteins, 9 (39%) were regulated by BMAL1 in at least one tissue and the abundance of many was restored in LMRE mice. Coagulation cascade proteins were also regulated by BMAL1 in both tissues (Figure 6E). This group included fibrinogen alpha and gamma chains (FGA and FGG) and regulators of protease activity including serine protease inhibitor c 1 (SERPINC1, also known as antithrombin-3) (Figure 6F). These pathways are regulated by the serpin family of serine protease inhibitors [63; 64]. In addition to SERPINC1, five other serpin family members were upregulated in *Bmal1* KO livers (SERPIN -A6, -B6B, -A1E, -F1 and -B1A). Interleukins 6 and 8 (IL-6 and IL-8), and chemokine c-c motif ligand 2 (CCL2, also called MCP-1) are rhythmically released from human skeletal myotubes synchronized *in vitro* [65]. However, these proteins were not detected in our dataset, perhaps due to their relatively low abundance in muscle prior to stimulation by exercise or circulating immune cells [66].

The number of secreted proteins affected by loss of *Bmal1* was more extensive in liver than muscle. Liver secreted proteins are enzymes, plasma proteins (e.g., albumin), hemostasis factors, apolipoproteins, components of the extracellular matrix, growth factors, and hormones [59; 67]. They contribute substantially to the makeup of the serum proteome [68; 69]. Metabolic pathways were among the functional enrichments of liver *Bmal1*-dependent secreted proteins, specifically cholesterol and lipid metabolism (Table S2, Figures S6A and S6B). We observed that the abundance of hepatic lipase (LIPC), the primary enzyme that catalyzes the hydrolysis of triglycerides to diacylglycerol and free fatty acids, was upregulated 2-fold in KO, rescued to WT levels in LMRE, and remarkably stable under TRF (Figures S6B). LIPC can exist within the liver bound to local endothelial cells or enter the circulation in complex with lipoproteins. It plays an important role in setting organismal levels of high-density lipoprotein (HDL) and low-density lipoprotein (LDL) cholesterols [70; 71]. Upregulation of LIPC was accompanied by downregulation of the LDL receptor (LDLR) and its transcriptional regulator SREBP [46; 72] (Figures S6B and S3F). This regulation of lipoprotein machinery likely contributes to the lipid phenotypes observed in clock mutant mice [11; 44; 73-75]. Interestingly, TRF tended to normalize liver abundance of SREBF1 and LDLR to WT levels.

Particularly noteworthy changes in *Bmal1* KO liver were a decrease in fibroblast growth factor 1 (FGF1) (Figure 7A), a metabolic regulator, and an increase in adiponectin (ADIPOQ) (Figure S3C). A similar but non-significant trend was also observed for ADIPOQ in muscle (Figure S3C). Adiponectin is an endocrine hormone produced by adipose tissue that regulates glucose and fatty acid metabolism in target tissues, including liver [76]. Its activity is associated with the suppression of metabolic impairments related to type 2 diabetes and fatty liver disease [77; 78]. There is evidence that adiponectin levels are intertwined with feeding and fasting behavior [79], however we did not observe a substantial change in ADIPOQ levels from AL to TRF in any genotype, suggesting a more prominent regulation by *Bmal1*.

**Figure 7.**
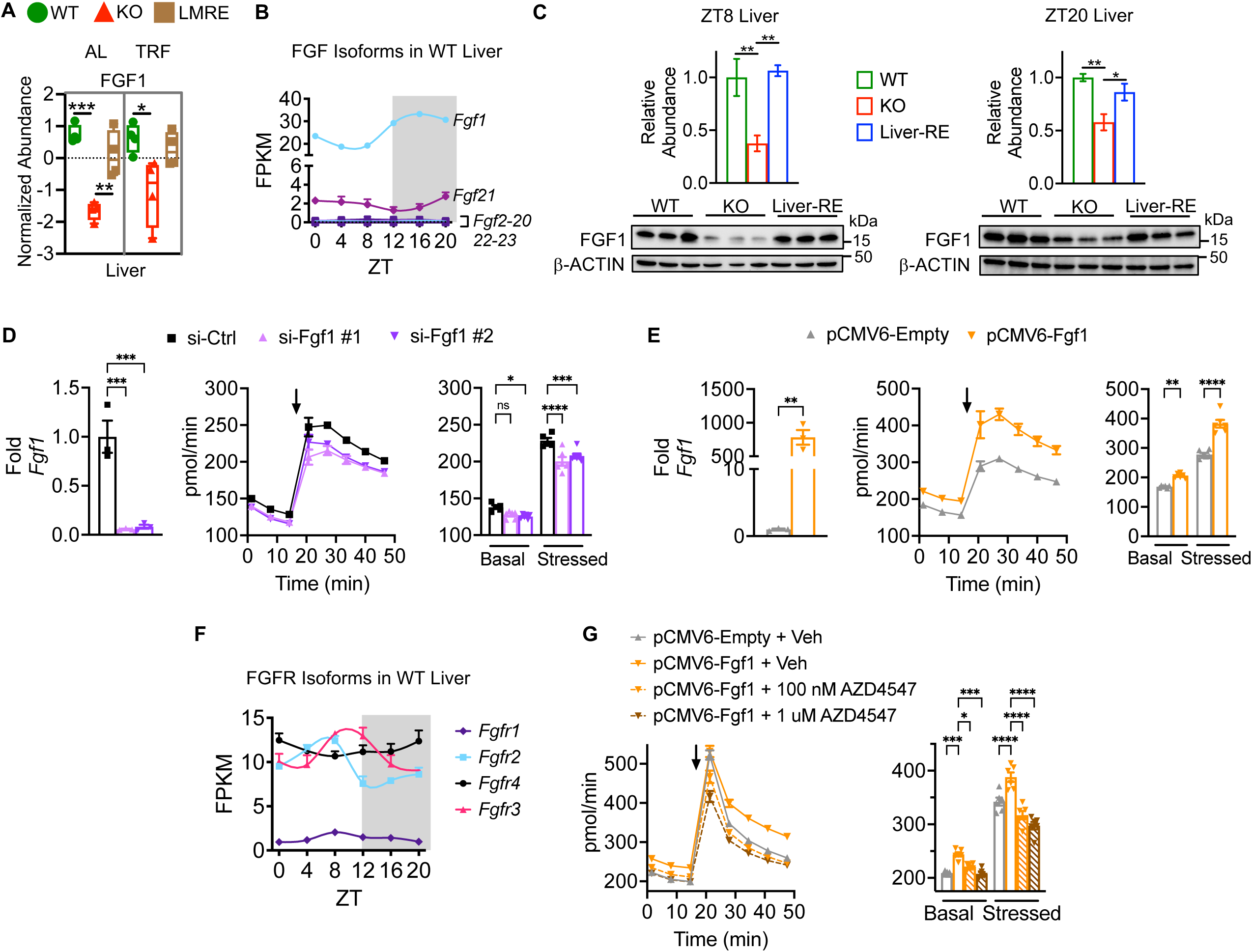
*Bmal1* regulates FGF1, which is necessary and sufficient for mitochondrial respiration in hepatocytes. (A) Abundance of fibroblast growth factor 1 (FGF1) protein in liver. AL – *ad libitum* feeding; TRF – time-restricted feeding. Two-way ANOVA with Tukey’s multiple comparisons test, * p<0.05, ** p<0.01, *** p<0.001, **** p<0.0001, n=4. (B) mRNA levels of FGF isoforms in WT (Alfp- and Hsa-Cre+) liver, n=3. ZT – zeitgeber time; ZT0 = lights on, ZT12 = lights off. (C) Western blot for FGF1 in liver at the indicated diurnal time points. Biological replicates are shown (bottom) and quantified (top). One-way ANOVA with Fisher’s LSD test, *p<0.05, ** p<0.01, n=3. Liver-RE = *Bmal1^stopFL/stopFL^*; *Alfp*-Cre^tg/0^, hepatocyte-specific reconstitution of *Bmal1*. (D and E) Knockdown or overexpression of *Fgf1* in AML12 hepatocytes. si-Ctrl = scrambled sequence; si-Fgf1 #1 and #2 both target *Fgf1* but by different sequences. pCMV6-Empty = control vector without *Fgf1* open reading frame clone. Left, qPCR, Student’s t-test, ** p<0.01, *** p<0.001, n=3. Middle, oxygen consumption rate in AML12 hepatocytes. Arrow indicates application of 0.5 uM carbonyl cyanide-p-trifluoromethoxyphenylhydrazone (FCCP) and 1 uM oligomycin. Right, quantification of middle. Basal and stressed values are averages of data points before and after drug application, respectively. Two-way ANOVA with Dunnett’s post-hoc test, * p<0.05, ** p<0.01, *** p<0.001, **** p<0.0001, ns – not significant, n=5-6. (F) mRNA levels of Fgf receptor (FGFR) isoforms in WT (Alfp- and Hsa-Cre+) liver. (G) Measurement of oxygen consumption rate as in D and E. AML12 cells were treated for 24 h with vehicle control (Veh) or the indicated concentration of the pan FGFR inhibitor AZD4547. Two-way ANOVA with Tukey’s post-hoc test, * p<0.05, *** p<0.001, **** p<0.0001, ns – not significant, n=5-6.

### 3.8. Bmal1 regulates FGF1 in liver

FGF1 (also known as acidic FGF) was originally identified as an endothelial growth factor (i.e., mitogen) but has reemerged as a metabolic regulator [80]. When challenged with a western-style diet or genetically-induced obesity, mice injected with recombinant FGF1 protein exhibit substantial rescue of hepatic steatosis and hepatic insulin sensitivity as well as suppression of hepatic glucose production [81; 82]. Despite these remarkable effects of FGF1 signaling on liver metabolism, the functional significance of the endogenous FGF1 protein in liver remains obscure.

Of the 22 FGF proteins, only FGF1 was quantified in our liver dataset, suggesting it is the most highly expressed isoform (Figure 7A). Indeed, *Fgf1* exhibited the highest expression at the RNA level, with 10-fold higher expression than *Fgf21* (Figure 7B). In addition, *Fgf1* exhibited small amplitude circadian variation, with peak expression during nighttime. Next, we confirmed *Bmal1*-dependent regulation of FGF1 by Western blot. FGF1 was significantly reduced in KO compared to WT livers harvested either during daytime (ZT8) or nighttime (ZT20), and rescue of only hepatocyte *Bmal1* (Liver-RE mice) was sufficient to restore liver FGF1 abundance (Figure 7C). Although *Fgf1* mRNA displayed a peak during the dark phase, we did not detect any changes at the protein level in a two time point analysis (Figure S7A). Because rescue of hepatocyte *Bmal1* rescued FGF1 abundance, our data indicate that hepatocytes are the main producers of FGF1 protein in liver. Additionally, AML12 cells, a non-transformed hepatocyte cell line, expressed FGF1 protein (Figure S7B).

### 3.9. Fgf1 Is necessary and sufficient for mitochondrial respiration in hepatocytes

A common thread among the reported effects of FGF1 on liver is mitochondrial function. We hypothesized that FGF1 could have a substantial impact on metabolism through the regulation of mitochondrial respiration, a process critical for catabolic and anabolic pathways in hepatocytes. To test this idea, we knocked down or overexpressed *Fgf1* in AML12 hepatocytes by siRNA and a mammalian expression vector, respectively. We confirmed efficient knockdown and overexpression by qPCR (Figures 7D and 7E) and then measured oxygen consumption, a readout of mitochondrial respiration, under basal or stressed conditions. Stressed conditions were created by adding the uncoupling agent FCCP and the ATP synthase inhibitor oligomycin. Because FGF1 regulates the growth of cells, we plated confluent cells and transfected them at the time of plating (reverse transfection). With this design, the number of cells was constant across the groups and thus did not confound measurements. We found that *Fgf1* knockdown lowered oxygen consumption and conversely that overexpression increased oxygen consumption (Figures 7D and 7E). Overexpression also increased extracellular acidification rate (a readout of glycolysis), yet to a lesser extent, and resulted in a larger change in oxygen consumption from baseline to stressed conditions, indicating a greater ability to meet metabolic demand (Figures S7C and S7D). These data show that FGF1 is necessary and sufficient for mitochondrial respiration in hepatocytes.

FGF1 exerts its effects through the FGF receptor tyrosine kinases (FGFRs). Treatment of hepatocytes with recombinant FGF1 protein reduced palmitate-induced lipid droplet formation in hepatocytes in an *Fgfr4*-dependent manner [81], however other *Fgfr* isoforms are also expressed in liver [83]. We assessed the expression of each *Fgfr* in WT liver and observed that *Fgfr2*, *Fgfr3*, and *Fgfr4* mRNAs were expressed similarly while *Fgfr1* expression was extremely low (Figure 7F). *Fgfr2* and *Fgfr3* exhibited small amplitude circadian variation, peaking near the end of the light phase, and *Fgfr4* was stable over the daily cycle. To determine if the observed effect of FGF1 on mitochondrial respiration depends on activation of its receptor, we treated cells with the pan-FGFR inhibitor AZD4547 [84]. At doses higher than ∼150 nM, AZD4547 inhibits all *Fgfr* isoforms [84]. Using either 100 nM or 1 uM, AZD4547 treatment was sufficient to block the increase in oxygen consumption rate and extracellular acidification rate induced by *Fgf1* overexpression (Figure 7G). Together, these data suggest that FGF1 promotes mitochondrial respiration in hepatocytes through an autocrine signaling mechanism. We surmise that the ability of FGF1 signaling to promote mitochondrial respiration depends on BMAL1 and contributes to the beneficial metabolic effects of FGF1 on liver metabolism [81; 82].

Due to the high affinity for heparan sulfate proteoglycans, which are components of the extracellular matrix that facilitate binding to FGF receptors, it is theorized that most FGF1 does not enter the circulation but rather signals in an autocrine or paracrine fashion [80]. Only FGF15/19, FGF21 and FGF23 are recognized as bona fide endocrine FGFs because they can bypass this mechanism. However, FGF1 is reported to circulate in mouse and human serum [85; 86]. Using a commercially available ELISA we detected FGF1 in WT serum at 0.988 ng/mL (Figure S7E). Considering that FGF1 is expressed in many organs of adult mice (Figure S7F), we sought to test whether it can be secreted from liver using an *ex vivo* approach. Briefly, an intact liver lobe was harvested, washed in PBS several times, and incubated for 1 h in serum-free media bubbled with carbogen. The media was then concentrated with a centrifugal filter and probed via Western blot. The media was positive for FGF1, suggesting FGF1 can be secreted from the liver (Figure S7G). Therefore, an investigation into a potential endocrine role of liver derived FGF1 is also warranted.

## 4. CONCLUSIONS

Through bulk proteomics, we find that organism-wide loss of *Bmal1* impacts a greater fraction of the liver proteome than skeletal muscle proteome. In line with our previous findings at the transcriptomic level [12], we also find that rescue of local liver and muscle clocks leads to a greater restoration of protein abundances in liver than in muscle, suggesting that the muscle clock relies more on external signals. However, our analysis in KO and local clock-restored mice under time-restricted feeding failed to yield appreciable changes in muscle, suggesting that inputs other than feeding-fasting may be important to support muscle protein abundances during the dark phase. Whilst we provide dark phase information, many proteins have recently been reported to exhibit circadian variation (reviewed in [87]) and therefore we expect additional layers of complexity to be revealed once this is taken into account.

We acknowledge that the bulk nature of the performed proteomics also limits sensitivity for detecting low or transiently expressed proteins. Therefore, cell-type-specific labeling and enrichment approaches [61; 88], or size-dependent fractionation, may lead to deeper proteome coverage. However, we were able to successfully reveal a role for local circadian clocks in liver and muscle in supporting the expression of secreted proteins. As part of these analyses, we identified a function of liver FGF1 in supporting mitochondrial respiration in hepatocytes via autocrine signaling. Future studies elucidating the local and distal targets of clock-dependent secreted proteins hold promise for identifying therapeutic targets for diverse metabolic diseases associated with dysfunction of liver and skeletal muscle, such as sarcopenia, NAFLD, and hepatocellular carcinoma.

## SUPPLEMENTAL FIGURE LEGENDS

## SUPPLEMENTAL TABLES

**Table S1. Proteome datasets**.

**Table S2. Pathway enrichments**.

**Table S3. *dryR* analysis**.

## ACKNOWLEDGEMENTS

This paper is dedicated to Paolo Sassone-Corsi, a truly inspiring scientist who continues to influence our work. We would also like to thank all members of the Parker and Koronowski laboratories for their efforts.

---

**Figure S1.**
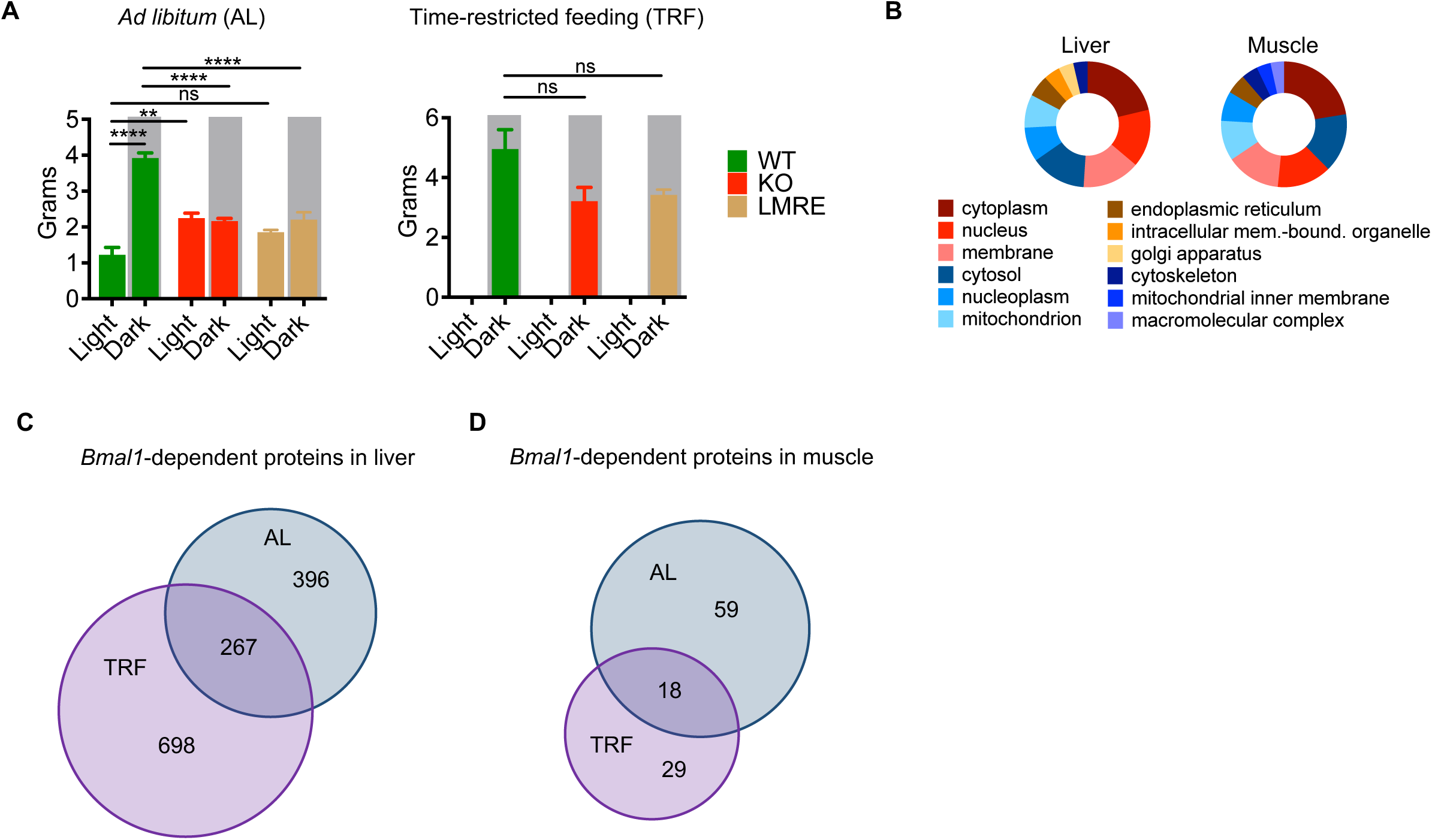
Analysis of food intake and quantified proteins. Related to Figure 1. (A) Average daily food intake. Two-way ANOVA with Tukey’s post-hoc test, ** p<0.01, **** p<0.0001, ns – not significant, n=3-5, LMRE n=2. (B) Gene ontology analysis of cellular compartment, showing the top localizations of quantified proteins from proteomic analysis. (C and D) Venn diagram analysis of proteins showing a significant difference in WT vs Bmal1-KO livers (C) or muscle (D) under AL or TRF.

**Figure S2.**
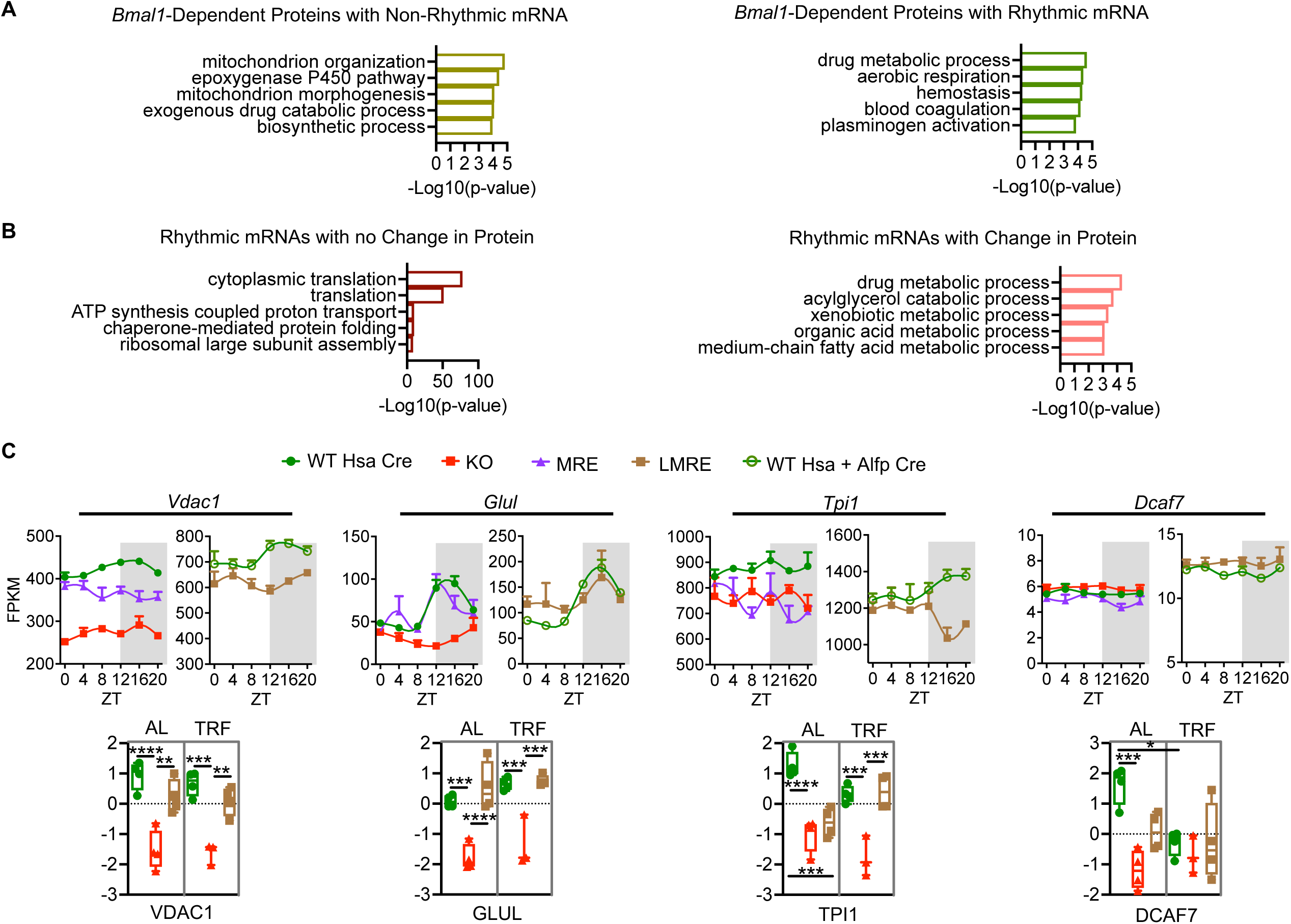
Analysis of the *Bmal1*-dependent proteome. Related to Figure 2. (A and B) Gene ontology enrichment (biological process) analysis showing the top 5 uniquely enriched pathways for each category of proteins. (C) Example genes/proteins in muscle. MRE = muscle specific *Bmal1* reconstitution. Voltage-dependent anion channel (VDAC1) and glutamate-ammonia ligase (GLUL) have rhythmic mRNA, lower daily average mRNA levels in KO, and a corresponding change in protein abundance. Triosephosphate isomerase 1 (TPI1) has non-rhythmic mRNA, lower daily average mRNA levels in KO, and a corresponding change in protein abundance. DDB1 and CUL4 associated factor 7 (DCAF7) protein abundances are regulated by *Bmal1* in a post-transcriptional manner. AL – *ad libitum* feeding; TRF – time-restricted feeding.

**Figure S3.**
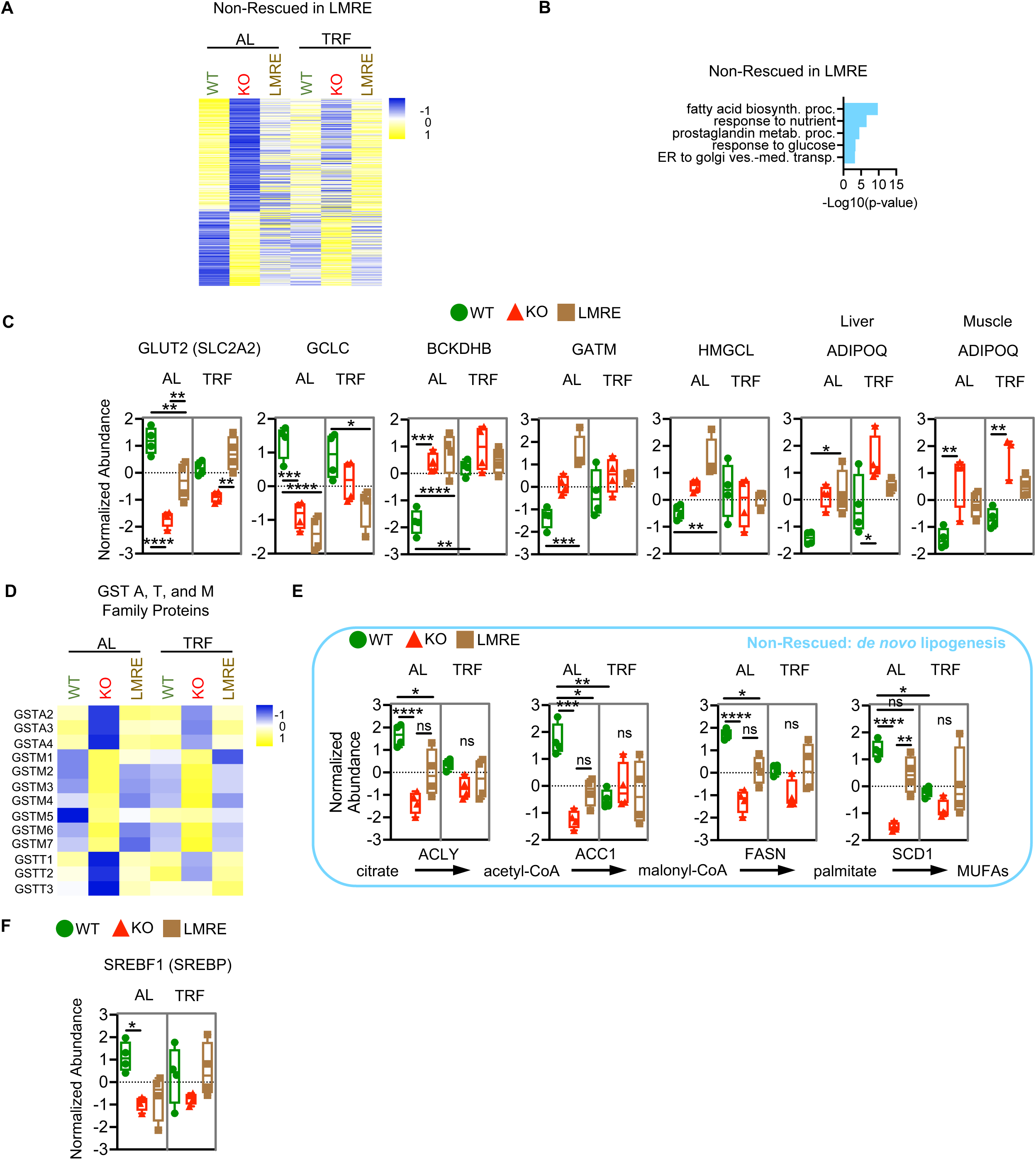
Analysis of liver proteins. Related to Figure 3. (A) Heatmaps of non-rescued proteins in liver. Non-rescued in LMRE = WT vs KO, q<0.05; LMRE vs KO q>0.05; WT vs LMRE q>0.05. (B) Gene ontology of non-rescued proteins in liver. (C) Examples of non-rescued proteins in liver (and muscle where indicated). Two-way ANOVA with Tukey’s multiple comparisons test, * p<0.05, ** p<0.01, *** p<0.001, **** p<0.0001, n=3-4. GLUT2 – facilitated glucose transporter, member 2; GCLC – glutamate-cysteine ligase, catalytic subunit; BCKDHB – branched chain ketoacid dehydrogenase E1, beta polypeptide; GATM – glycine amidinotransferase; HMGCL – hydroxymethylglutaryl-CoA lyase, mitochondrial; ADIPOQ – adiponectin. AL – *ad libitum* feeding; TRF – time-restricted feeding. (D) Heatmap of quantified glutathione *S*-transferases (GSTs) of the theta (T), alpha (A), and mu (M) classes. (E) Protein abundances of proteins involved in *de novo* lipogenesis in liver. ACLY – ATP citrate lyase; ACC1 (ACACA) – acetyl-coenzyme A carboxylase alpha; FASN – fatty acid synthase; SCD1 – stearoyl-coenzyme A desaturase 1. Statistics same as in C.

**Figure S4.**
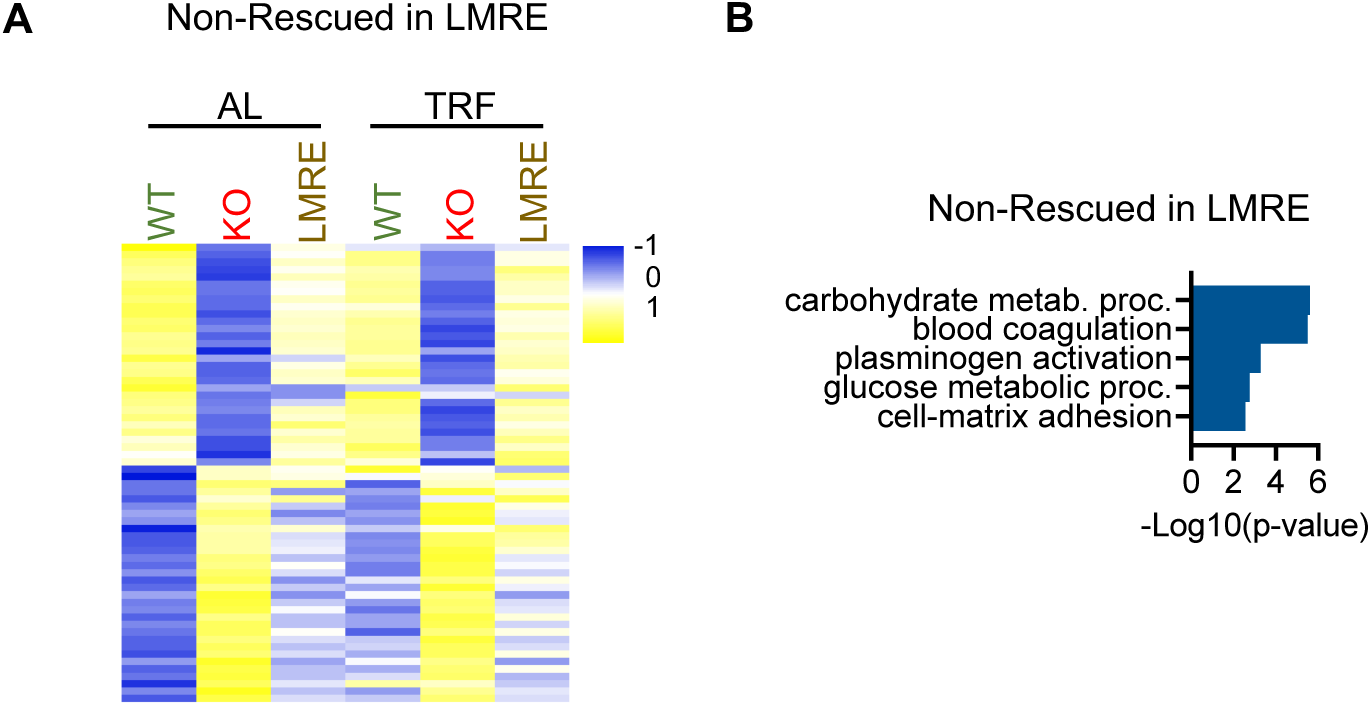
Analysis of muscle proteins. Related to Figure 4. (A) Heatmaps of non-rescued proteins in muscle. Non-rescued in LMRE = WT vs KO, q<0.05; LMRE vs KO q>0.05; WT vs LMRE q>0.05. (B) Gene ontology of non-rescued proteins in muscle.

**Figure S5.**
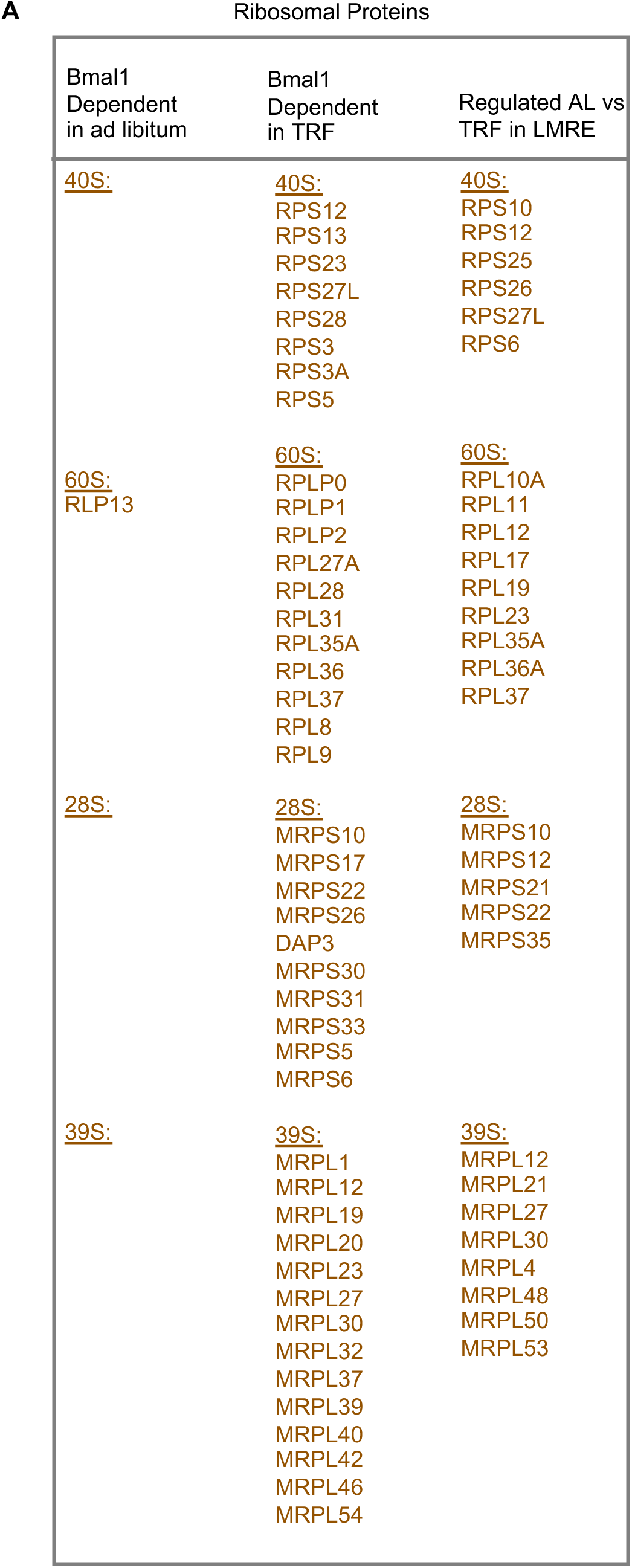
Analysis of ribosomal proteins in liver. Related to Figure 5. (A) List of quantified ribosomal proteins in liver. Student’s t-test with false discovery rate correction, q<0.05, n=4. *Bmal1*-dependent in ad libitum = WT AL vs KO AL. *Bmal1*-dependent in TRF = WT TRF vs KO TRF. Feeding-dependent = AL vs TRF in LMRE.

**Figure S6.**
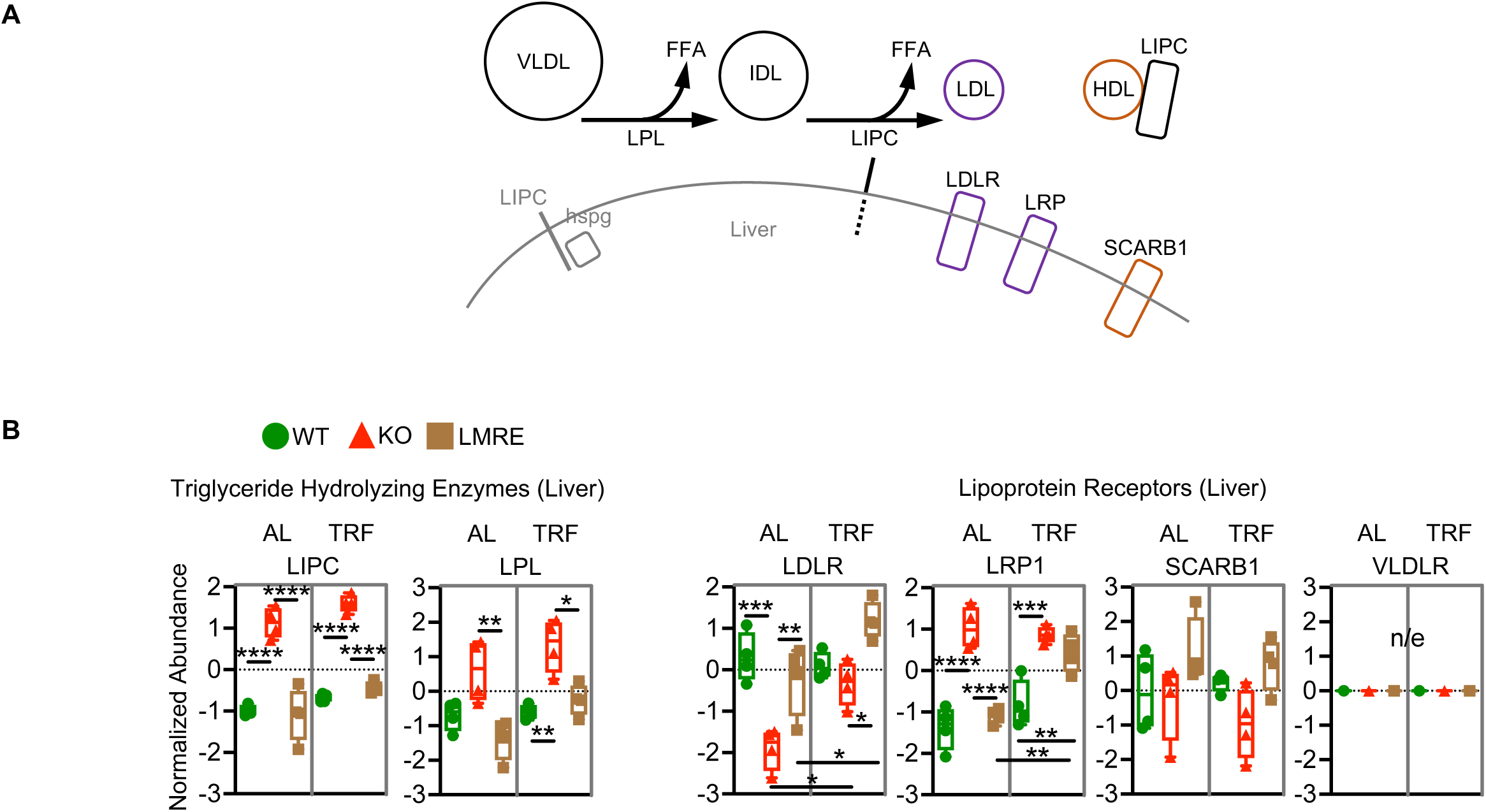
Analysis of secreted proteins. Related to Figure 6. (A) Simplified scheme of systemic triglyceride metabolism involving the liver. VLDL – very-low-density lipoprotein; IDL – intermediate-density lipoprotein; LDL – low-density lipoprotein; HDL – high-density lipoprotein; FFA – free fatty acid; LPL – lipoprotein lipase; LIPC – lipase, hepatic; hspg – heparan sulfate proteoglycans; LDLR – low density lipoprotein receptor; LRP – low density lipoprotein receptor-related protein 1; SCARB1 – scavenger receptor class B, member 1. (B) Abundances of proteins shown in A. Two-way ANOVA with Tukey’s multiple comparisons test, * p<0.05, ** p<0.01, *** p<0.001, **** p<0.0001, n=4. n/e – not expressed in liver.

**Figure S7.**
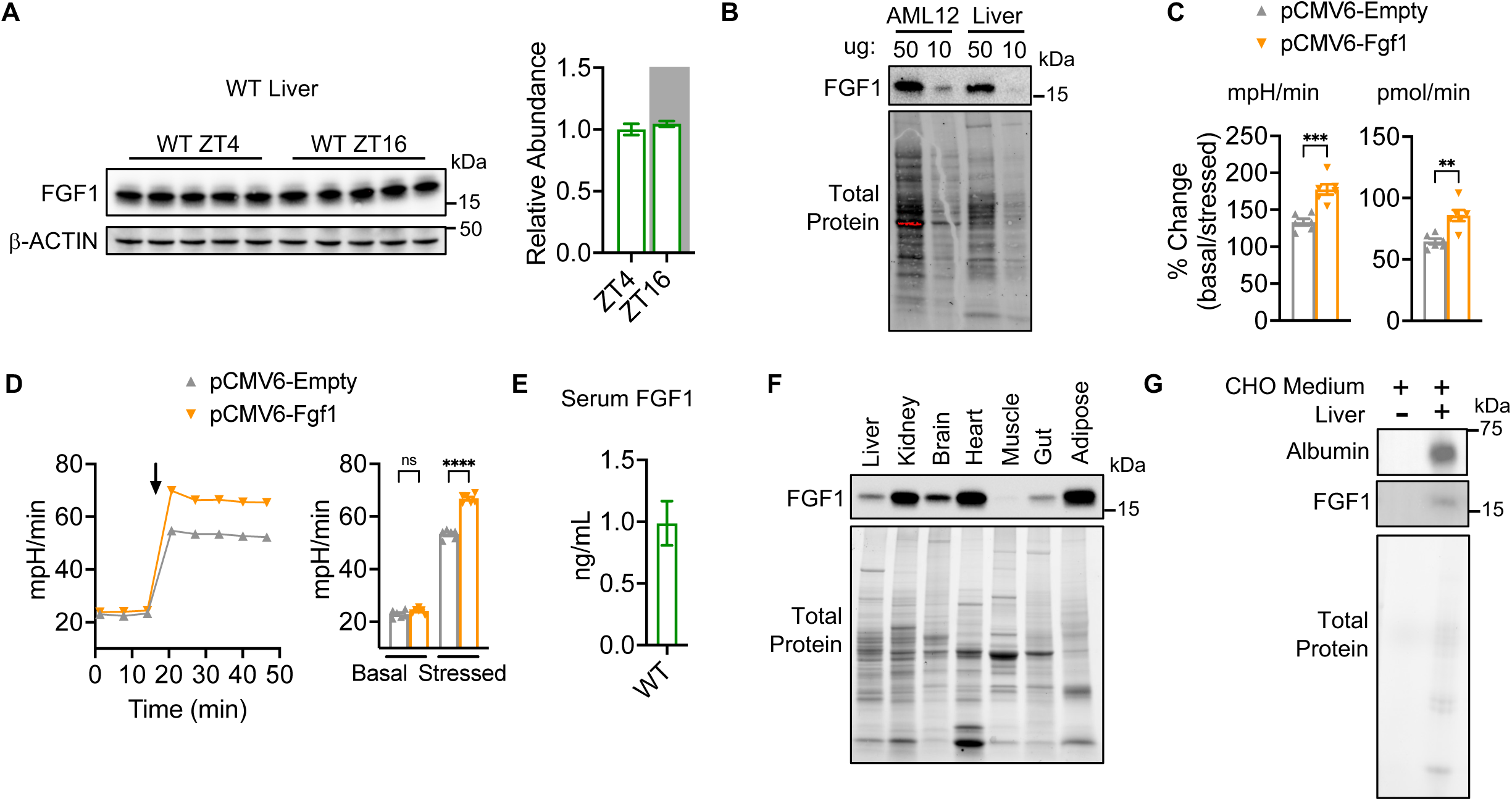
Analysis of FGF1. Related to Figure 7. (A) Western blot analysis of whole cell lysates from liver harvested at the indicated diurnal time points. ZT – zeitgeber time; ZT0 = lights on, ZT12 = lights off. Right, quantification of n=5 biological replicates. (B) Example western blot showing protein expression of FGF1 in AML12 hepatocytes and liver whole cell lysate. Total protein from gel. Ug - indicates the amount of protein loaded. (C) Change in the rate (left, extracellular acidification; right, oxygen consumption) from basal to stressed conditions in AML12 hepatocytes with control transfection or overexpression of *Fgf1*. Student’s t-test, ** p<0.01, *** p<0.001, n=5-6. (D) Left, extracellular acidification rate in AML12 hepatocytes with control transfection or overexpression of *Fgf1*. Arrow indicates application of 0.5 uM carbonyl cyanide-p-trifluoromethoxyphenylhydrazone (FCCP) and 1 uM oligomycin. Right, quantification of left. Basal and stressed values are averages of data points before and after drug application, respectively. Two-way ANOVA with Sidak’s post-hoc test, **** p<0.0001, ns – not significant, n=5-6. (E) Quantification of FGF1 protein in WT serum by ELISA. (F) Example western blot showing the relative expression of FGF1 across metabolic tissues. Total protein from gel. Muscle is gastrocnemius. Brain is cortex. (G) Example western blot showing the presence of FGF1 in chemically-defined media (bubbled with carbogen) incubated *ex vivo* for 45 min with a whole liver lobe. Albumin is a known liver-secreted protein and is included as a positive control. Total protein from gel.

